# Dietary soy shapes murine microbiota to consolidate the mucosal IgA response through T follicular helper cells

**DOI:** 10.1101/2025.08.12.669459

**Authors:** Kisara Hattori-Muroi, Hikari Maruta, Daisuke Takahashi, Yusuke Kinashi, Kouya Hattori, Yuma Kabumoto, Yumiko Fujimura, Satoshi Tsukamoto, Koichiro Suzuki, Hiroyuki Oguchi, Yumi Ogihara, Yotaro Kodaira, Emi Hayashi, Kokona Takano, Seiga Komiyama, Naoki Morita, Hanako Naganawa-Asaoka, Yuki Oya, Yuka Saito, Wakana Ohhashi, Shunsuke Kimura, Reiko Shinkura, Tsukasa Matsuda, Koji Hase

## Abstract

The commensal microbiota plays a crucial role in shaping mucosal immunity, particularly in the induction of T follicular helper (Tfh) cells and IgA production. Here, we demonstrate that dietary soy elicits a robust Tfh cell and IgA response in Peyer’s patches of weaning mice. Soy feeding promotes the expansion of two principal commensal bacterial species, *Limosilactobacillus reuteri* and *Muribaculum intestinale*. Mechanistically, *L. reuteri* provides cognate antigens for Tfh cell activation, while *M. intestinale* functions as an adjuvant by promoting IL-1β production from myeloid cells. The resulting IgA exhibits polyreactivity and enhances protection against *Salmonella* infection. These findings highlight the specific interplay among dietary components, intestinal microbiota, and mucosal immunity, thereby establishing a diet-microbe-immune axis that shapes host defense in early life. This axis represents a promising therapeutic target for developing novel strategies to enhance resistance to enteric pathogens.

**Graphical abstract:** 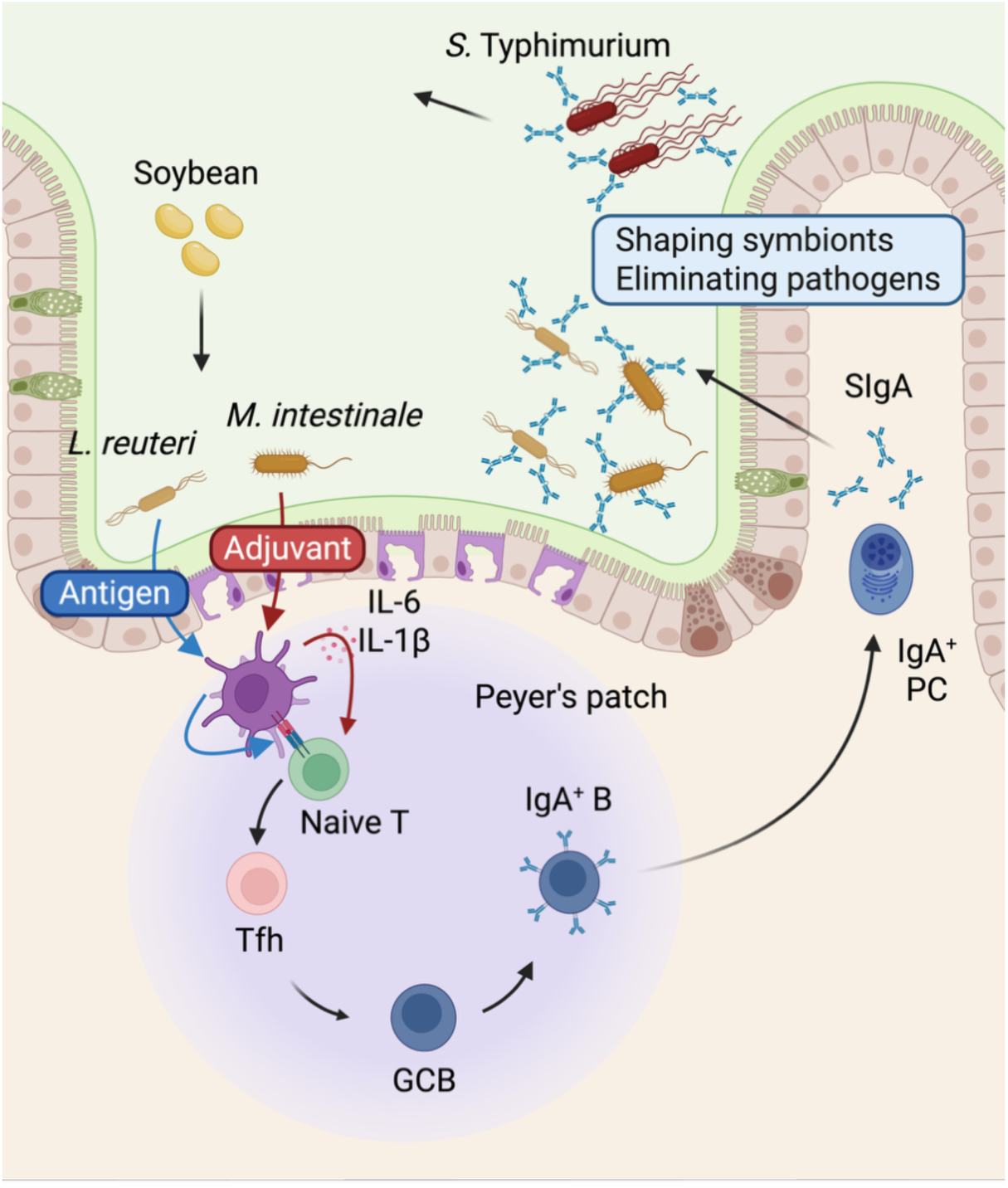

**Highlights:**

- Dietary soy promotes T follicular helper (Tfh) cell and IgA responses in the Peyer’s patches of weaning mice.
- Soy enhances the colonization of two key commensals*, Limosilactobacillus reuteri* and *Muribaculum intestinale,* in the small intestine.
- These two bacteria synergize to drive mucosal immunity: *L. reuteri* provides cognate antigens, while *M. intestinale* provides an adjuvant signal by promoting IL-1β production from myeloid cells.
- Soy-induced IgA is polyreactive and protects against *Salmonella* infection.

## Introduction

The human intestinal tract is home to an estimated 40 trillion commensal bacteria.^1^ These bacteria play a pivotal role in establishing the intestinal immune system, the body’s largest immune system, and maintaining a symbiotic relationship with the host. The gut-associated lymphatic tissues (GALTs), which develop within the intestinal tract, are primarily responsible for gut mucosal immune responses, with a particular emphasis on the production of immunoglobulin A (IgA).^2^ The interaction of polymeric IgA with a specific antibody transporter, termed the polymeric Ig receptor, enables the transcytosis of polymeric IgA across intestinal epithelial cells. This process culminates in the accumulation of secretory IgA (SIgA).^3^ SIgA serves as a frontline barrier on mucosal surfaces by neutralizing toxins and eliminating pathogenic bacteria.^4^ Several mechanisms have been identified for the elimination of pathogenic bacteria by SIgA. These include inhibiting proliferation, reducing motility through binding to flagella, and clumping or chaining of bacteria, particularly pathogens such as *Salmonella* spp., *Vibrio cholerae*, and *Enterococcus faecalis*, to exclude them from the gut.^5,6^ Interestingly, SIgA can also bind to beneficial commensal bacteria in the gut, aiding their adhesion to the mucosal surface and contributing to the symbiont inhabitance.^7,8^ This multifaceted role of SIgA is indispensable in maintaining a balance in the gut microbiota and preserving a symbiotic relationship. Notably, IgA deficiency causes an increased systemic translocation of specific taxa, such as Enterobacteriaceae, leading to aberrant activation of the systemic immune response to the translocated bacteria.^9^ This highlights the significance of mucosal IgA in maintaining systemic immune homeostasis by preventing microbial translocation.

The induction of IgA occurs via two distinct pathways, T cell-independent (TI) and T cell-dependent (TD).^10^ The TI pathway does not induce IgA affinity maturation through somatic hypermutation. Instead, it produces IgA that recognizes a broad range of commensal bacteria.^11–13^ This polyreactivity allows TI IgA to bind a broad range of antigens, providing a generalized barrier in the gut. On the other hand, the TD pathway involves the activation of B cells in the GALT, which generates hypermutated IgA that targets specific commensal bacteria with high affinity. Importantly, this high-affinity IgA can also exhibit polyreactivity when there are shared epitopes among different commensal bacteria.^14^ Thus, high-affinity polyreactive IgA contributes to maintaining the balance in the symbiotic commensal microbiota by affecting their fitness, motility, and metabolic functions.^15–17^ For instance, activation-induced cytidine deaminase (AID)-deficient mice lacking IgA, as well as AID^G23S^ mutant mice lacking only high-affinity IgA, exhibit commensal dysbiosis.^18,19^ Moreover, the high-affinity IgA primarily recognizes highly penetrant commensal pathobionts and also provides protection against pathogens. Such a dual role of high-affinity IgA underscores its importance in both maintaining gut health and protecting against disease.^14,20^

T follicular helper (Tfh) cells, a specialized subset of CD4^+^ T cells, play a crucial role in producing high-affinity antibodies, including IgA, in the secondary lymphoid tissues.^21^ In the systemic immune system, Tfh cells transiently increase in numbers during bacterial or viral infections, forming germinal centers (GC) in collaboration with antigen-specific B cells to promote the production of antigen-specific IgG. Conversely, in the Peyer’s patches (PPs), one of the GALT that serve as the major inductive site for mucosal immune responses, a substantial population of Tfh cells is present even under steady-state conditions, continuously regulating TD IgA production.^22^ The differentiation of Tfh cells primarily begins when dendritic cells (DCs) in the PPs capture antigens and present them to naïve CD4^+^ T cells. This process involves stimulation through the T cell receptor (TCR) by the peptide:MHCII complex and by co-stimulatory factors, with interleukin-6 (IL-6) playing a particularly significant role in Tfh cell differentiation, including induction of the lineage-defining transcription factor Bcl-6 in mice.^23^ Pre-Tfh cells encountering cognate B cells at the T-B border undergo maturation through interactions with B cells, further upregulating the expression of C-X-C motif chemokine receptor 5 (CXCR5), allowing their localization within the GC.^24^ IL-1β produced by CX3CR1^+^ monocytes acts as a key signal that, in combination with IL-6, drives the differentiation of Tfh cells.^25^ Conversely, IL-2 acts antagonistically on the Tfh cell differentiation program. IL-2 high producers after TCR stimulation give rise to Tfh cells, in other words, a subset of naïve T cells that received the strongest TCR signals is likely to differentiate into Tfh cells.^26–28^ Activated B cells receive CD40 stimulation from Tfh cells and cytokine stimulation, particularly IL-21, leading to class switching from IgM^+^ B cells to IgA^+^ B cells.^29,30^ Subsequently, B cells engage with FDCs to capture antigens and undergo clonal expansion and somatic hypermutation (SHM).^31^ SHM introduces random mutations and clones with increased affinity to the antigen are selected by Tfh cells. Iterations of SHM and selection by Tfh cells, a process known as affinity maturation, enhances antibody affinity.^32,33^ Dysfunction of Tfh cells leads to incomplete induction of affinity maturation, resulting in the generation of low-affinity IgA against bacteria, ultimately causing dysbiosis and impaired barrier function.^34^ In support of these phenomena, it has recently been shown that high-affinity IgA produced in a TD manner is polyreactive to multiple enteric bacteria.^14,35^ The differentiation of effector T cells is affected by specific gut bacteria. T helper 17 (Th17) cells and regulatory T (Treg) cells, both of which are abundant in the lamina propria (LP), are decreased in germ-free mice, with several inducers, including gut bacteria and their metabolites, having been identified.^36–39^ Tfh cells also require gut microbiota for their differentiation in PPs.^40^ By contrast, GCB cells remain unchanged or even increase in mesenteric lymph nodes compared to specific pathogen-free (SPF) mice.^41^ Nevertheless, antigen-specific B cells from GCs in germ-free (GF) mice exit at a faster rate compared to SPF mice, and the released IgA exhibits extremely low affinity for any antigens, namely, dietary antigens, self-antigens, and gut bacterial antigens, despite undergoing SHM to some extent. This suggests that an adequate selection of IgA^+^ B cells by Tfh cells is critical for the production of high-affinity IgA. Tfh cells in germ-free mice can be partially restored through the administration of Toll-like receptor (TLR) ligands.^40^ However, the number of PP Tfh cells is not influenced just by the amount of commensal microbiota but rather by its composition. In particular, segmented filamentous bacteria (SFB), a pathobiont in the mouse ileum, is known as a potent inducer of PP Tfh cell differentiation, which can lead to autoantibody-mediated arthritis.^42^ Although SFB is almost the only species among commensal bacteria that has been clearly shown to induce the differentiation of Tfh cells, SFB is absent in human microbiota. Nevertheless, a substantial number of Tfh cells consistently exist in the human PPs, indicating that environmental factors other than SFB also contribute to Tfh cell differentiation.

Given that factors other than SFB must contribute to Tfh cell differentiation, particularly in humans, we explored how environmental factors shape this process. We focused on dietary interventions as a key investigative strategy, given that diet plays a critical role in shaping the gut microbiota. Notably, the development of the gut microbiota begins in tandem with the introduction of solid foods during the weaning period,^43^ making this a critical window for establishing mucosal immunity. In this study, we demonstrate that dietary soy is a key factor that promotes Tfh cell development and subsequent IgA responses in PPs. This effect is driven by a specific synergistic interaction between two commensals enriched by soy: *Limosilactobacillus reuteri*, which provides cognate antigens, and *Muribaculum intestinale*, which functions as an adjuvant by promoting IL-1β production from myeloid cells. This diet-microbe axis culminates in the production of high-affinity, polyreactive IgA capable of protecting the host from enteric pathogens. Our study thus reveals a specific diet-microbe-immune axis that shapes mucosal immunity in early life.

## Results

### Semi-purified diet feeding reduces the number of Tfh, GCB, and IgA^+^ B cells in PPs

To modify the composition of gut microbiota through dietary intervention, we opted for either the unpurified diet CE-2 or the semi-purified diet AIN-93G (Table S1). The CE-2 diet, composed of crude materials derived from plants, yeasts, and animals, provides abundant substrates that intestinal microbiota can utilize. In contrast, AIN-93G, primarily composed of purified materials, allows for rapid absorption by the host and offers fewer microbiota-accessible substrates. After feeding weaning mice with either CE-2 or AIN-93G for four weeks, we analyzed the gut microbiota in the distal small intestine and Tfh cells in PP. As we described in an earlier study,^44^ the β-diversities of the gut microbiota were significantly different between the two groups (Figure 1A). Additionally, α-diversity decreased in the AIN-93G-fed group compared to the CE-2-fed group (Figure 1B). At the family level, the AIN-93G group exhibited a significant decrease in *Muribaculaceae* (formerly known as family S24-7) and a corresponding increase in *Erysiperotrichaceae* and *Peptostreptococcaceae*, which became the most abundant family (Figure 1C). The total bacterial count in the distal small intestine also decreased in the AIN-93G group (Figure S1A). Importantly, CXCR5^+^Bcl-6^+^ Tfh cells in the distal PPs of the AIN-93G group were reduced by one-third in frequency and one-fifth in cell number compared to the CE-2 group (Figures 1D and 1E). Bcl-6^+^GL^+^ GCB cells did not differ significantly in percentage, whereas the cell number was reduced to about one-third in the distal PPs in the AIN-93G group (Figures 1F and 1G). A similar reduction was observed in the number of IgA^+^ B cells (Figures 1H and 1I). The changes in Tfh cells, GCB, and IgA^+^ B cells in PPs were limited to the distal small intestine and were not observed in the proximal small intestine (Figure 1D-1I), which has a lower bacterial burden, suggesting that these changes occur in a gut microbiota-dependent fashion. In line with these alterations in the inductive sites, we found that IgA-producing plasma cells (IgA⁺IRF4⁺) were significantly reduced in the lamina propria (LP) of both the proximal and distal small intestine in the AIN-93G group (Figures 1J and 1K). Consistent with the decreased number of plasma cells, the concentration of IgA in feces was less than half in the AIN-93G group compared to the CE-2 group (Figure 1L). Additionally, a decrease in Th17 cells was also observed in the LP of the distal small intestine in the AIN-93G group (Figures S1B and S1C).

**Figure 1.**
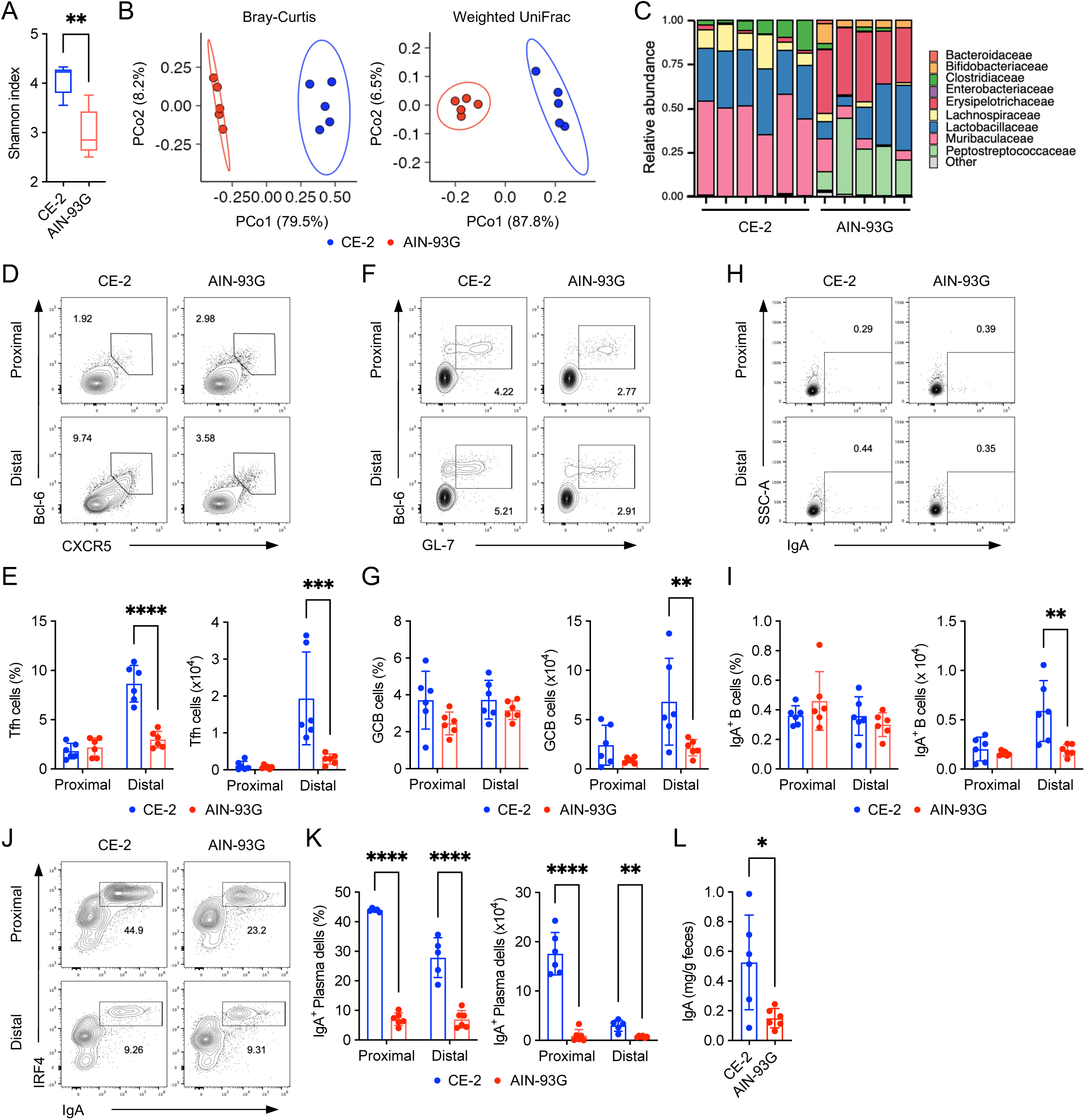
A semi-purified diet reduces the number of Tfh, GCB, and IgA⁺ B cells in PPs. (A-C) Analysis of the distal small intestinal microbiota in mice fed either the CE-2 or the AIN-93G diet for 4 weeks starting from 3 weeks of age (*n* = 5-6 mice/group). (A) Species richness (Shannon entropy). (B) Principal coordinate analysis (PCoA) of Bray-Curtis and weighted UniFrac distances. (C) Microbiota composition at the family level. (D-K) Flow cytometry analysis of lymphocytes in Peyer’s patches (PPs) and lamina propria (LP) from mice fed either the CE-2 or the AIN-93G diets for 4 weeks starting from 3 weeks of age. (D and E) Frequency and total number of CXCR5⁺Bcl-6⁺ Tfh cells in proximal and distal PPs. Representative plots of CXCR5 and Bcl-6 staining within the CD4⁺TCRβ⁺Foxp3⁻CD25⁻ gate are shown in (D) (*n* = 6 mice/group, mean ± s.d.). (F and G) Frequency and total number of GL-7⁺Bcl-6⁺ GCB cells in proximal and distal PPs. Representative plots of GL-7 and Bcl-6 staining within the CD19⁺ gate are shown in (F) (*n* = 6 mice/group, mean ± s.d.). (H and I) Frequency and total number of IgA⁺ B cells in proximal and distal PPs. Representative plots of IgA staining within the CD19⁺ gate are shown in (H) (*n* = 6 mice/group, mean ± s.d.). (J and K) Frequency and total number of IRF4⁺IgA⁺ plasma cells in proximal and distal LP. Representative plots of IgA and IRF4 staining within the CD19⁻ gate are shown in (J) (*n* = 5-6 mice/group, mean ± s.d.). (L) Concentration of IgA in fecal samples from mice described in (D-K), measured by ELISA (*n* = 6 mice/group, mean ± s.d.). Data are representative of at least two independent experiments. Statistical analysis was performed by (A) Mann-Whitney test, (L) Welch’s t-test, or (E, G, I, and K) Two-way ANOVA followed by Šídák’s multiple comparisons test. \**p* < 0.05; \*\**p* < 0.01; \*\*\**p* < 0.001; \*\*\*\**p* < 0.0001. See also Figure S1 amd S2.

### Soy supplementation to the AIN-93G diet restores the PP Tfh cells and subsequent IgA responses

To investigate which ingredient in the CE-2 diet contributes to Tfh cell induction, we supplemented the AIN-93G diet with one of five dietary ingredients (i.e., soy, germ oil, whole wheat, manganese, and iron) that are either unique to the CE-2 diet or abundant in it. Weaning mice were maintained on these modified AIN-93G diets for four weeks. Only soy supplementation restored both the frequency and number of Tfh cells in the distal PPs to a level comparable to that in the CE-2 diet group (Figures 2A and 2B). Similarly, PP GCB and IgA^+^ B cells in the distal PPs significantly increased in the soy-supplemented AIN-93G diet (AIN-93G + Soy) group compared to the AIN-93G diet group (Figures 2C, 2D, S2A, and S2B). Consistent with this, soy supplementation increased IgA-producing plasma cells in the LP of both proximal and distal small intestine (Figures 2E and 2F), leading to a higher IgA concentration (Figure 2G). Notably, these effects were commensal bacteria-dependent, as the AIN-93G + Soy diet failed to increase any of the PP Tfh, GCB, and IgA^+^ B cells in the distal PPs of GF mice compared to the AIN-93G group (Figures S2C-S2I). Furthermore, continuous intake of soy was essential for maintaining these immune responses. Switching from the AIN-93G + Soy diet to the AIN-93G diet after two weeks reduced Peyer’s patch Tfh cells to levels observed in mice maintained on the AIN-93G diet throughout (Figures S3A-S3C). The frequencies of GCB and IgA⁺ B cells also returned to baseline levels, although their cell numbers remained slightly higher (Figures S3D-S3G).

**Figure 2.**
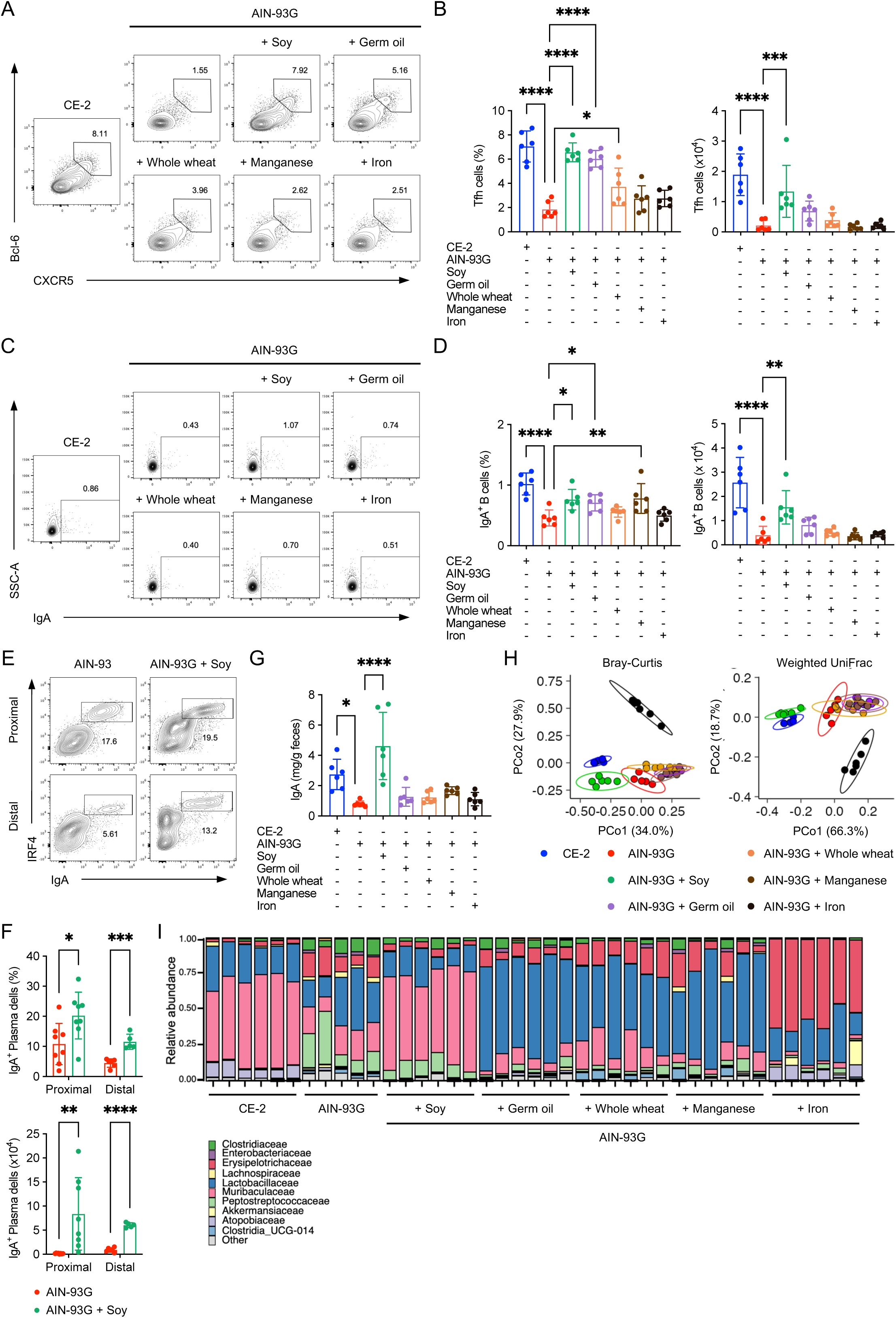
Soy supplementation to the AIN-93G diet restores the PP Tfh cell population and subsequent IgA responses. (A-F) Flow cytometry analysis of lymphocytes in PPs and LP from mice fed CE-2, AIN-93G, or AIN-93G diets supplemented with representative CE-2 ingredients for 4 weeks starting from 3 weeks of age. (A and B) Frequency and total number of CXCR5⁺Bcl-6⁺ Tfh cells in distal PPs. Representative plots of CXCR5 and Bcl-6 staining within the CD4⁺TCRβ⁺Foxp3⁻CD25⁻ gate are shown in (A) (*n* = 6 mice/group, mean ± s.d.). (C and D) Frequency and total number of IgA⁺ B cells in distal PPs. Representative plots of IgA staining within the CD19⁺ gate are shown in (C) (*n* = 6 mice/group, mean ± s.d.). (E and F) Frequency and total number of IRF4⁺IgA⁺ plasma cells in proximal and distal LP. Representative plots of IgA and IRF4 staining within the CD19⁻ gate are shown in (E) (*n* = 5-8 mice/group, mean ± s.d.). (G) Concentration of IgA in fecal samples from mice described in (A-F), measured by ELISA (*n* = 6 mice/group, mean ± s.d.). (H and I) Analysis of the distal small intestinal microbiota from mice fed CE-2 diet, AIN-93G diet, or AIN-93G + Soy diets for 4 weeks starting from 3 weeks of age (*n* = 5-6 mice/group). (H) PCoA of Bray-Curtis and weighted UniFrac distances. (I) Microbiota composition at the family level. Data are representative of at least two independent experiments. Statistical analysis was performed by (B, D, G) One-way ANOVA followed by Dunnett’s or Tukey’s multiple comparisons test, or (F) Two-way ANOVA followed by Šídák’s multiple comparisons test. \**p* < 0.05; \*\**p* < 0.01; \*\*\**p* < 0.001; \*\*\*\**p* < 0.0001. See also Figures S2-S4.

To confirm the essential role of Tfh cells in this process, we utilized mice lacking Tfh cells (*Cd4*-cre;*Bcl6*^f/f^). In these mice, the AIN-93G + Soy diet feeding failed to induce both GCB and IgA⁺ B cells in the distal PPs (Figures S4A-S4F). Furthermore, the induction of Tfh, GCB, and IgA⁺ B cells by the AIN-93G + Soy diet was diminished in M cell-deficient mice (*Vil1*-cre;*Tnfrsf11a*^f/f^), suggesting that M cell–mediated translocation of luminal antigens is required for this germinal center response (Figures S4G-S4L). In addition to Tfh cell induction, similar to the CE-2 diet, AIN-93G + Soy diet feeding also increased Th17 cells in distal small intestinal LP (Figures S4M and S4N).

We next examined the impact of soy supplementation on the gut microbiota. Although the total amount of bacteria in the distal small intestine tended to increase in the AIN-93G + Soy diet group compared to the AIN-93G diet group, it remained lower compared to the CE-2 diet group (Figure S4O). However, the α-diversity of commensal microbiota, which decreased in the AIN-93G diet group compared to the CE-2 diet group, was also restored in the AIN-93G + Soy diet group (Figure S4P). Consequently, the composition of the distal small intestinal microbiota of the AIN-93G + Soy diet group closely resembled that of the CE-2 diet group (Figures 2H and 2I). As we and others previously reported, AIN-93G diet feeding led to the decolonization of SFB,^44,45^ which was not prevented by soy supplementation (Figure S4Q). Together, these data demonstrate that among the CE-2 ingredients, soy is responsible for inducing PP Tfh cells and the subsequent IgA response by altering the composition, rather than the quantity, of small intestinal microbiota.

### Soy diet feeding augments specific bacterial clades that induce PP Tfh cells

Given the dynamic changes in the microbiota composition between the AIN-93G and the AIN-93G + Soy diet groups, we next sought to identify the specific bacteria responsible for Tfh cell induction. To narrow down candidate bacteria, we analyzed Tfh cells under two experimental conditions: antibiotic treatment and variations in breeding facility environments. First, weaning mice were administered vancomycin (VCM), streptomycin (SM), or erythromycin (EM) via the drinking water for four weeks, concurrent with feeding of either an AIN-93G diet or an isocaloric AIN-93G + Soy diet (Table S2). As anticipated, each antibiotic treatment group exhibited a distinct microbiota composition, reflecting the differences in the antibiotic spectrum (Figure 3A). The total number of bacteria was extremely low in the EM-treated group, whereas the reduction was moderate in the VCM and SM-treated groups (Figure S5A). The induction of Tfh cells in the distal PPs of the AIN-93G + Soy diet group was lost in all antibiotic-treated groups (Figures 3B and 3C). Likewise, all the antibiotic treatments eliminated the increase in GCB and IgA^+^ B cells caused by soy supplementation (Figures S5B-5E).

**Figure 3.**
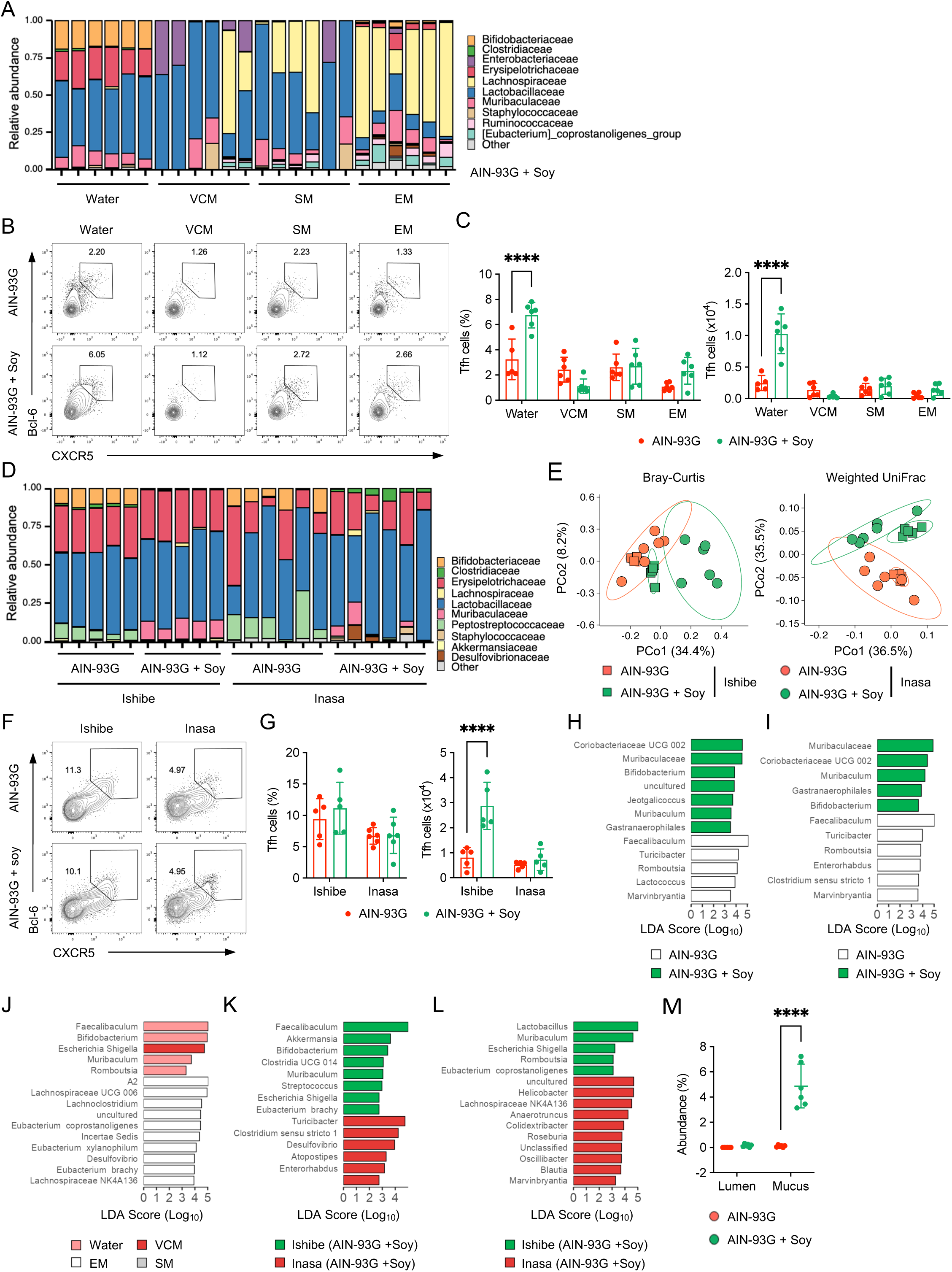
Soy diet feeding augments specific bacterial clades that induce PP Tfh cells. (A) Composition of the distal small intestinal microbiota at the family level in mice fed either the AIN-93G or the AIN-93G + Soy diet and treated with antibiotics (vancomycin, VCM; streptomycin, SM; or erythromycin, EM) for 4 weeks starting from 3 weeks of age (*n* = 6 mice/group). (B and C) Flow cytometry analysis of CXCR5⁺Bcl-6⁺ Tfh cells in the distal PPs of mice treated with antibiotics or water for 4 weeks. Representative plots of CXCR5 and Bcl-6 staining within the CD4⁺TCRβ⁺Foxp3⁻CD25⁻ gate are shown in (B), and quantification of frequency and total number is shown in (C) (*n* = 6 mice/group, mean ± s.d.). (D and E) Microbiota analysis in mice from different breeding facilities (Ishibe or Inasa) fed either the AIN-93G or the AIN-93G + Soy diet for 4 weeks. (D) Microbiota composition at the family level. (E) PCoA of Bray-Curtis and weighted UniFrac distances (*n* = 5-6 mice/group). (F and G) Flow cytometry analysis of CXCR5⁺Bcl-6⁺ Tfh cells in the distal PPs of mice from Ishibe and Inasa facilities. Representative plots of CXCR5 and Bcl-6 staining within the CD4⁺TCRβ⁺Foxp3⁻CD25⁻ gate are shown in (F), and quantification of frequency and total number is shown in (G) (*n* = 5-6 mice/group, mean ± s.d.). (H-M) Linear discriminant analysis effect size (LEfSe) of bacterial communities. (H and I) Comparison between AIN-93G and AIN-93G + Soy diets in distal small intestinal contents (H) and mucus (I) of mice from Ishibe facility. (J) Comparison between untreated and antibiotic-treated mice in the intestinal contents of mice from the Ishibe facility. (K and L) Comparison between Ishibe and Inasa facilities in small intestinal contents (K) and mucus (L). (M) Abundance of *M. intestinale* in intestinal contents and mucus of mice from the Ishibe facility (*n* = 6 mice/group). Data are representative of at least two independent experiments. Statistical analysis was performed by (C, G, and M) Two-way ANOVA followed by Šídák’s multiple comparisons test. \*\*\*\**p* < 0.0001. See also Figures S5-S7.

In a subsequent experiment, we fed the AIN-93G + Soy diet to the mice born in either the Ishibe facility of CLEA Japan, which we routinely use, or the Inasa facility of Japan SLC. The mice from the Inasa facility exhibited a distinct microbial composition compared to those from the Ishibe facility after being fed the AIN-93G + Soy diet (Figures 3D and 3E). Unlike the observations in Ishibe mice, the soy diet did not impact the abundance of Tfh, GCB, and IgA^+^B cells in distal PPs of Inasa mice (Figures 3F, 3G, and S6A-S6D). These results suggest that a specific bacterial clade in Ishibe mice mediates the induction of PP Tfh cells and the subsequent IgA^+^ B cell response to the AIN-93G + Soy diet. To identify such a candidate, we therefore used the linear discriminant analysis (LDA) effect size (LEfSe) method^46^ to compare the bacterial communities in distal small intestinal content and mucus across the three independent comparisons: AIN-93G vs AIN-93G + Soy diet feeding (Figures 3H and 3I), untreated vs antibiotic-treated (Figure 3J), and Ishibe vs Inasa mice fed the AIN-93G + Soy diet (Figures 3K and 3L). The results revealed a consistent enrichment of a specific genus, *Muribaculum*, in the groups that exhibited an increase in Tfh cells across all three comparisons (Figures 3H-3L). Alignment of the 16S rRNA gene sequence via nucleotide BLAST demonstrated that this genus corresponds to *Muribaculum intestinale*, and shotgun metagenome sequencing confirmed that this species was specifically enriched in the AIN-93G + Soy diet group (Figure S6E). Interestingly, 16S rRNA gene analysis revealed that while *M. intestinale* was present in both the luminal content and the mucus-associated fraction, its relative abundance was consistently higher in the mucus layer, suggesting a preference for this niche (Figure 3M).

### A combination of soy components is required for the growth of *M. intestinale* and subsequent Tfh cell induction

We subsequently investigated how soy feeding facilitates the growth of *M. intestinale*. Because soy oil is a primary lipid source in the AIN-93G diet (Table S2), we focused on other major components of soy, such as soy protein and carbohydrates, including fibers. We therefore prepared an AIN-93G diet supplemented with soy protein and/or soy fiber (Table S3) and fed it to weaning mice for four weeks. However, the abundance of *M. intestinale* in the distal small intestine was not affected by the individual or combined supplementation of soy protein and fiber in the AIN-93G diet (Figure S7A). Consistent with this, frequency and number of Tfh cells in the distal PPs were not changed by feeding the soy protein and fiber-supplemented AIN-93G diet (Figures S7B and S7C).

Considering that *M. intestinale* has a diverse repertoire of glycoside hydrolases, including α-galactosidase,^47^ we next evaluated the effect of oligosaccharides, namely stachyose and raffinose, which are abundant in soy.^48^ We, therefore, prepared an AIN-93G diet supplemented with raffinose and stachyose in addition to soy protein and soy fiber (AIN-93G + SP/SF/Ra/St) (Table S3). Four weeks of this diet feeding resulted in the colonization of *M. intestinale* to levels comparable to that of the AIN-93G + Soy diet group in both the distal small intestinal content and the mucus layer (Figures S7D and S7E). Nevertheless, the AIN-93G + SP/SF/Ra/St diet had no effect on the abundance of Tfh cells in the distal PPs, suggesting that colonization by *M. intestinale* alone is insufficient to induce Tfh cells (Figures S7F and S7G). This finding prompted us to compare the microbial compositions between AIN-93G + SP/SF/Ra/St and AIN-93G + Soy diet groups by shotgun metagenome sequencing. While the intake of the AIN-93G + SP/SF/Ra/St diet substantially increased the abundance of *M. intestinale*, that of *Lactobacillus* spp. was much lower compared to the AIN-93G + Soy diet group (Figure S7H). In particular, *Limosiactobacillus reuteri* (formerly known as *Lactobacillus reuteri*) was nearly eliminated in the AIN-93G + SP/SF/Ra/St diet group.

### Co-colonization of *L. reuteri* and *M. intestinale* synergistically induces the PP Tfh cells and subsequent IgA responses

To investigate the individual and synergistic effects of *L. reuteri* and *M. intestinale* on the induction of Tfh, GCB, and IgA⁺ B cells in the distal PPs, we established gnotobiotic mice associated with either *L. reuteri*, *M. intestinale*, or both. Their successful colonization in the lumen and mucus layer of the distal small intestine was confirmed by qPCR (Figures S8A and S8B). These gnotobiotic mice were fed the AIN-93G + Soy diet for four weeks, which did not alter the total bacterial load (Figure S8C). A small amount of bacterial DNA was detected in the germ-free group, likely originating from nonviable *Lactococcus lactis* present in the casein.^49^ Interestingly, the frequency of Tfh cells in the distal PPs increased specifically in mice co-colonized with both *L. reuteri* and *M. intestinale* (Figures 4A and 4B). The total number of Tfh cells increased in mice mono-colonized with *M. intestinale*, irrespective of *L. reuteri* co-colonization (Figure 4B). Although mono-colonization with either *L. reuteri* or *M. intestinale* induced GCB cells, co-colonization with both species led to the greatest increase in GCB cells (Figures 4C and 4D). Accordingly, significant increases in IgA⁺ B cells and fecal IgA levels were observed only under co-colonization conditions (Figures 4E-4G).

**Figure 4.**
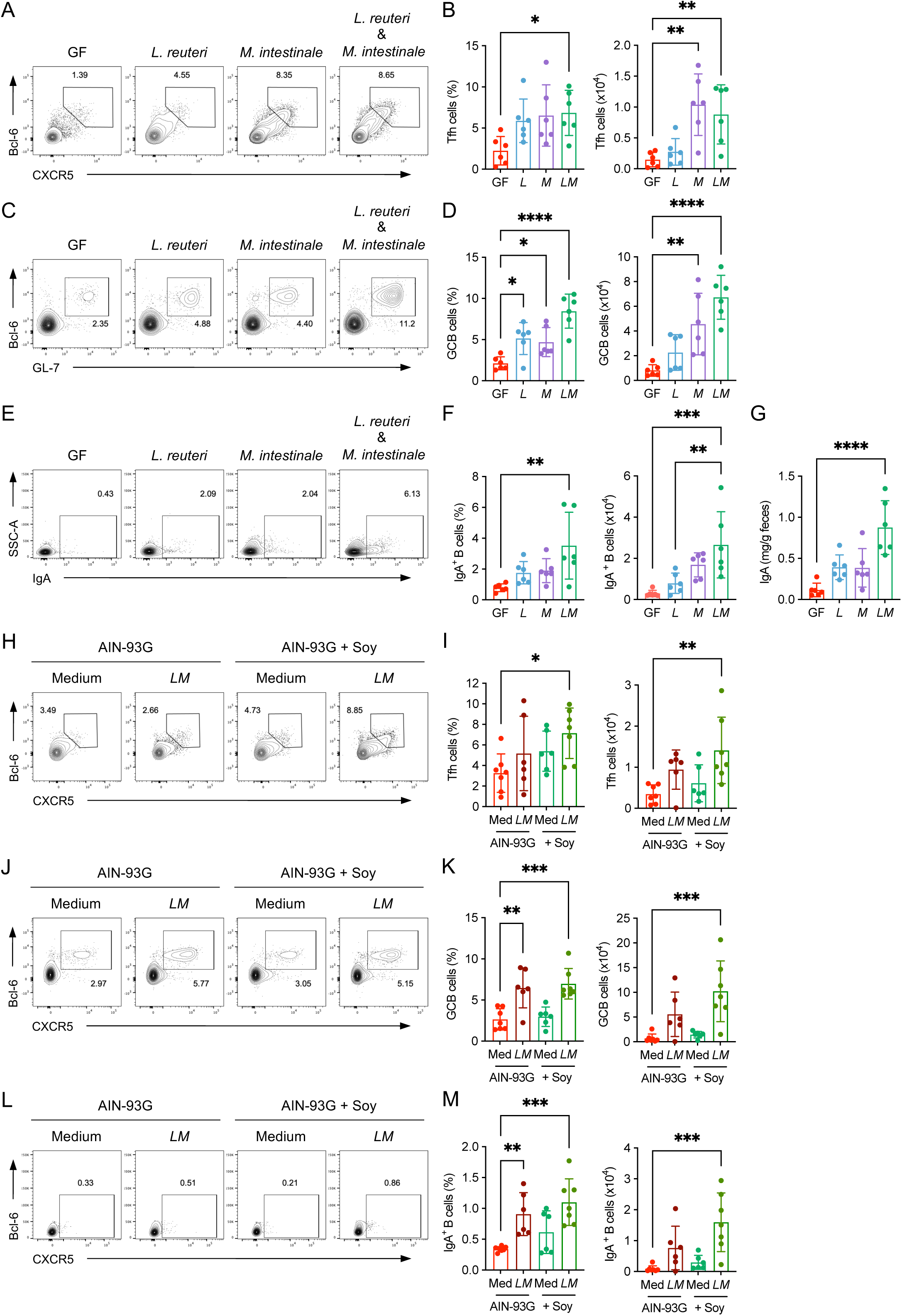
Co-colonization of L. reuteri and M. intestinale synergistically induces PP Tfh, GCB, and IgA⁺ B cells. (A-F) Flow cytometry analysis of lymphocytes in distal PPs from GF, *L. reuteri* (*L*), *M. intestinale* (*M*) mono-colonized, and *L. reuteri* and *M. intestinale* (*LM*) co-colonized gnotobiotic mice fed the AIN-93G + Soy diet for 4 weeks starting from 3 weeks of age. (A and B) Frequency and total number of CXCR5⁺Bcl-6⁺ Tfh cells in distal PPs. Representative plots of CXCR5 and Bcl-6 staining within the CD4⁺TCRβ⁺Foxp3⁻CD25⁻ gate are shown in (A) (*n* = 6 mice/group, mean ± s.d.). (C and D) Frequency and total number of GL-7⁺Bcl-6⁺ GCB cells in distal PPs. Representative plots of GL-7 and Bcl-6 staining within the CD19⁺ gate are shown in (C) (*n* = 6 mice/group, mean ± s.d.). (E and F) Frequency and total number of IgA⁺ B cells in distal PPs. Representative plots of IgA staining within the CD19⁺ gate are shown in (E) (*n* = 6 mice/group, mean ± s.d.). (G) Concentration of IgA in fecal samples from mice described in (A-F), measured by ELISA (*n* = 6 mice/group, mean ± s.d.). (H-M) Flow cytometry analysis of lymphocytes in distal PPs from mice from the Inasa facility inoculated with *L. reuteri* and *M. intestinale* (*LM*) or control culture medium (Med) and fed either the AIN-93G or the AIN-93G + Soy diet for 4 weeks starting from 3 weeks of age. (H and I) Frequency and total number of CXCR5⁺Bcl-6⁺ Tfh cells in distal PPs. Representative plots of CXCR5 and Bcl-6 staining within the CD4⁺TCRβ⁺Foxp3⁻CD25⁻ gate are shown in (H) (*n* = 6 mice/group, mean ± s.d.). (J and K) Frequency and total number of GL-7⁺Bcl-6⁺ GCB cells in distal PPs. Representative plots of GL-7 and Bcl-6 staining within the CD19⁺ gate are shown in (J) (*n* = 6 mice/group, mean ± s.d.). (L and M) Frequency and total number of IgA⁺ B cells in distal PPs. Representative plots of IgA staining within the CD19⁺ gate are shown in (L) (*n* = 6 mice/group, mean ± s.d.). Data are representative of at least two independent experiments. Statistical analysis was performed by (B, D, F, and G) One-way ANOVA followed by Tukey’s multiple comparisons test, or (I, K, and M) Two-way ANOVA followed by Šídák’s multiple comparisons test. \*\**p* < 0.05; \*\**p* < 0.01; \*\*\**p* < 0.001; \*\*\*\**p* < 0.0001. See also Figures S8 and S9.

To validate these findings in a conventional setting, we performed a colonization experiment using mice from the Inasa facility, in which both *L. reuteri* and *M. intestinale* showed low abundance (Figures 3D, 3K, and 3L). While forced inoculation of these two bacteria failed to establish colonization in mice fed the AIN-93G diet, their colonization was successful in the distal small intestine of mice fed the AIN-93G + Soy diet (Figures S8D and S8E). Crucially, the induction of Tfh cells in the distal PPs of Inasa mice required the combination of bacterial inoculation and AIN-93G + Soy diet feeding; neither intervention alone was sufficient (Figures 4H and 4I). A similar synergistic effect was observed for the induction of GCB and IgA⁺ B cells in the distal PPs (Figures 4J-4M). Notably, even when the co-inoculation of the two bacterial strains failed to establish stable colonization upon the AIN-93G feeding, a modest increase in GCB and IgA⁺ B cells was still detected, suggesting that transient exposure to these bacteria may be sufficient to elicit a partial immune response (Figures 4J-4M).

Finally, to examine whether the synergy was specific to *M. intestinale*, we tested whether it could be substituted by another commensal bacterium, *Faecalibaculum rodentium*, whose abundance was higher in mice fed the AIN-93G diet (Figure S7H). We established gnotobiotic mice co-colonized with *L. reuteri* and *F. rodentium* and confirmed their successful colonization (Figures S9A-S9C). Unlike co-colonization with *M. intestinale*, co-colonization with *F. rodentium* did not increase Tfh cells in the distal PPs (Figures S9D and S9E). Although the frequency and number of GCB cells were increased in mice colonized with *L. reuteri* and *F. rodentium* compared with GF mice, the effect was significantly weaker than that observed with *L. reuteri* and *M. intestinale* (Figures S9F and S9G). Moreover, *L. reuteri* and *F. rodentium* co-colonization had no effect on IgA⁺ B cells in the distal PPs (Figures S9H and S9I).

These results collectively indicate that while *M. intestinale* alone can expand the number of Tfh cells, the synergistic action of *M. intestinale* and *L. reuteri* is required for fully activating germinal center reactions and expanding IgA⁺ B cell populations, ultimately leading to increased TD IgA production.

### *L. reuteri* provides antigens while *M. intestinale* promotes IL-1β production from DCs to cooperatively boost Tfh cell induction

To elucidate the mechanisms underlying Tfh cell induction by *L. reuteri* and *M. intestinale*, we first sought to investigate the significance of the bacterial antigens as TCR ligands. To this end, Tfh cells were sort-purified from the distal PPs of soy-fed *Bcl6*-tdTomato-cre^ERT2^ reporter mice, and fused with αβTCR^-/-^ BW5147 cells to establish Tfh hybridomas. Tfh hybridomas were co-cultured with bone marrow-derived dendritic cells (BMDCs) pre-pulsed with small intestinal content derived from the GF, *L. reuteri* or *M. intestinale* gnotobiotic mice. The reactivity of the Tfh hybridoma population was assessed by measuring IL-2 production in the culture supernatant. The small intestinal content from *L. reuteri* gnotobiotic mice induced higher IL-2 production compared to the content from GF mice, while that from *M. intestinale* gnotobiotic mice had little effect (Figure 5A). To further evaluate the reactivity of Tfh cells to *L. reuteri*, we established a total of 81 single Tfh hybridoma clones, each possessing a unique TCRα and β chain. Assessment of their reactivity by IL-2 production revealed that a substantial portion of the clones responded to *L. reuteri* lysates. Specifically, 2 out of 81 clones (2.5%) were strongly reactive, while 7 (8.6%) and 41 (50.6%) clones responded at medium and low levels, respectively; the remaining 31 clones (38.3%) showed no response (Figures 5B and S10A). Collectively, these results indicate that *L. reuteri*-derived antigens are recognized by a significant fraction of the PP Tfh cell population expanded by the soy diet.

**Figure 5.**
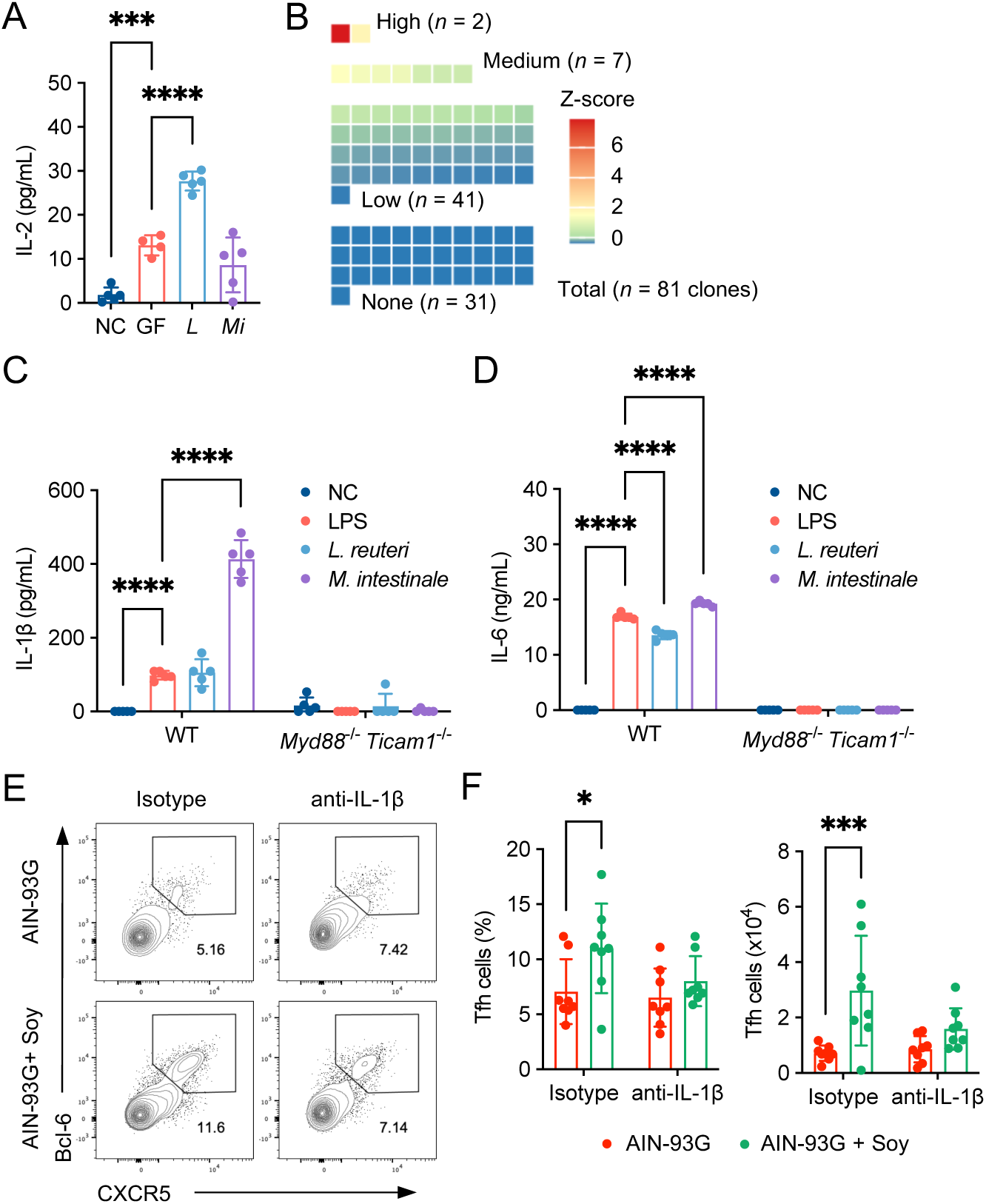
*L. reuteri* provides antigens while *M. intestinale* promotes IL-1β production from DCs to cooperatively boost Tfh cell induction. (A) IL-2 concentration in the supernatant of PP Tfh hybridoma pools co-cultured for 24 hours with BMDCs and lysates of small intestinal contents from GF, *L. reuteri* (*L*), or *M. intestinale* (*M*) gnotobiotic mice, or PBS (NC) (*n* = 5 technical replicates/group, mean ± s.d.). (B) Heat map showing Z-scores of IL-2 concentration in the supernatant of 81 single PP Tfh hybridoma clones, each possessing a unique TCRα and β pair, co-cultured with BMDCs and lysates of *L. reuteri*. The clones were categorized by their reactivity levels. (C and D) Concentration of IL-1β (C) and IL-6 (D) in the supernatant of BMDCs from wild-type (WT) or *Myd88*^-/-^*Ticam1*^-/-^ (MyD88 and TRIF-deficient) mice, stimulated with LPS, L. reuteri, or M. intestinale lysates, or unstimulated (NC) (*n* = 5 technical replicates/group, mean ± s.d.). (E and F) Flow cytometry analysis of CXCR5⁺Bcl-6⁺ Tfh cells in the distal PPs of mice fed either the AIN-93G or the AIN-93G + Soy diet and administered with an anti-IL-1β neutralizing antibody or isotype control for the last two weeks. Representative plots of CXCR5 and Bcl-6 staining within the CD4⁺TCRβ⁺Foxp3⁻CD25⁻ gate are shown in (E), and quantification of frequency and total number is shown in (F) (*n* = 8 mice/group, mean ± s.d.). Data in (A-D) are representative of two independent experiments. Data in (E and F) are from one representative experiment of at least two. Statistical analysis was performed by (A) One-way ANOVA followed by Tukey’s multiple comparisons test, or (C, D and F) Two-way ANOVA followed by Šídák’s multiple comparisons test. \**p* < 0.05; \*\*\**p* < 0.001; \*\*\*\**p* < 0.0001. See also Figure S10.

Given the established synergistic role of IL-6 and IL-1β in Tfh cell priming by myeloid cells,^24,25^ we performed single-cell RNA sequencing (scRNA-seq) on PP CD11c⁺ cells to identify myeloid subsets involved in Tfh cell differentiation. While cluster sizes were comparable between the AIN-93G and AIN-93G + Soy diet groups (Figure S10B), the proportion of cells expressing *Il1b* and its expression level were increased in the conventional DC1 (cDC1), cDC2, and monocyte-derived DC (moDC) clusters from the AIN-93G + Soy diet group (Figure S10C). In contrast, *Il6* expression was low across all CD11c⁺ subsets in both groups, at least at the transcriptome level (Figure S10C).

To further investigate the impact of *L. reuteri* and *M. intestinale* on cytokine production by DCs, we stimulated BMDCs with heat-killed bacteria or lipopolysaccharide (LPS) as a positive control. Intriguingly, *M. intestinale* significantly promoted both IL-1β and IL-6 production from BMDCs, with a more pronounced effect than *L. reuteri* (Figures 5C and 5D). The upregulation of these cytokines by *M. intestinale* was dependent on Toll-like receptor (TLR) signaling, as the effect was abolished in *Myd88*⁻/⁻*Trif*⁻/⁻ BMDCs (Figures 5C and 5D).^50^ Finally, we investigated the impact of IL-1β depletion on PP Tfh cell induction by the soy-commensal axis. Weaning mice fed either the AIN-93G or AIN-93G + Soy diet were administered an anti-IL-1β neutralizing antibody or an isotype control for two weeks. Neutralization of IL-1β abolished the Tfh cell-inducing effect of the AIN-93G + Soy diet (Figures 5E and 5F). Based on these observations, we conclude that *M. intestinale* facilitates the differentiation of PP Tfh cells by stimulating DCs to produce IL-1β through TLR activation.

### Soy-induced IgA is polyreactive and enhances resistance to *S*. Typhimurium infection

We subsequently assessed the features of lamina propria IgA induced by the AIN-93G + Soy diet. We sequenced the immunoglobulin heavy chain (IgH) genes from IgA⁺ plasma cells in the distal small intestine of mice fed either the AIN-93G or AIN-93G + Soy diet. Both groups had similar α-diversities in their IgA repertoires (Figure 6A and 6B). Approximately 90% of the sequences from AIN-93G and AIN-93G + Soy diet groups possessed mutations and a high ratio of replacement (R) to silent (S) mutations in complementarity-determining regions 1 to 3 (CDR1-3), compared with those in framework regions 1 to 3 (FWR1-3). Notably, the calculated affinity maturation index (R_CDR/_S_Total_+R_CDR_)^51^ was higher in the AIN-93G + Soy diet group, although the mutation rate at each region was comparable between the two groups (Figures 6C and S11A). In addition, AIN-93G + Soy diet feeding significantly increased the proportion of IgA-coated bacteria, especially IgA^high^ bacteria, in both proximal and distal small intestinal content but not in the colon content (Figure 6D, 6E, S11B and S11C). This suggests that soy supplementation enhances the selection of high-affinity IgA^+^ B cells by Tfh cells, resulting in increased production of IgA that targets bacterial antigens. Importantly, *L. reuteri* was highly coated by IgA only in the AIN-93-Soy diet group, while *M. intestinale* was not a direct target of IgA (Figures 6F and S11D), consistent with our finding that Tfh hybridomas generated from the AIN-93-Soy diet group primarily recognize *L. reuteri* antigens.

**Figure 6.**
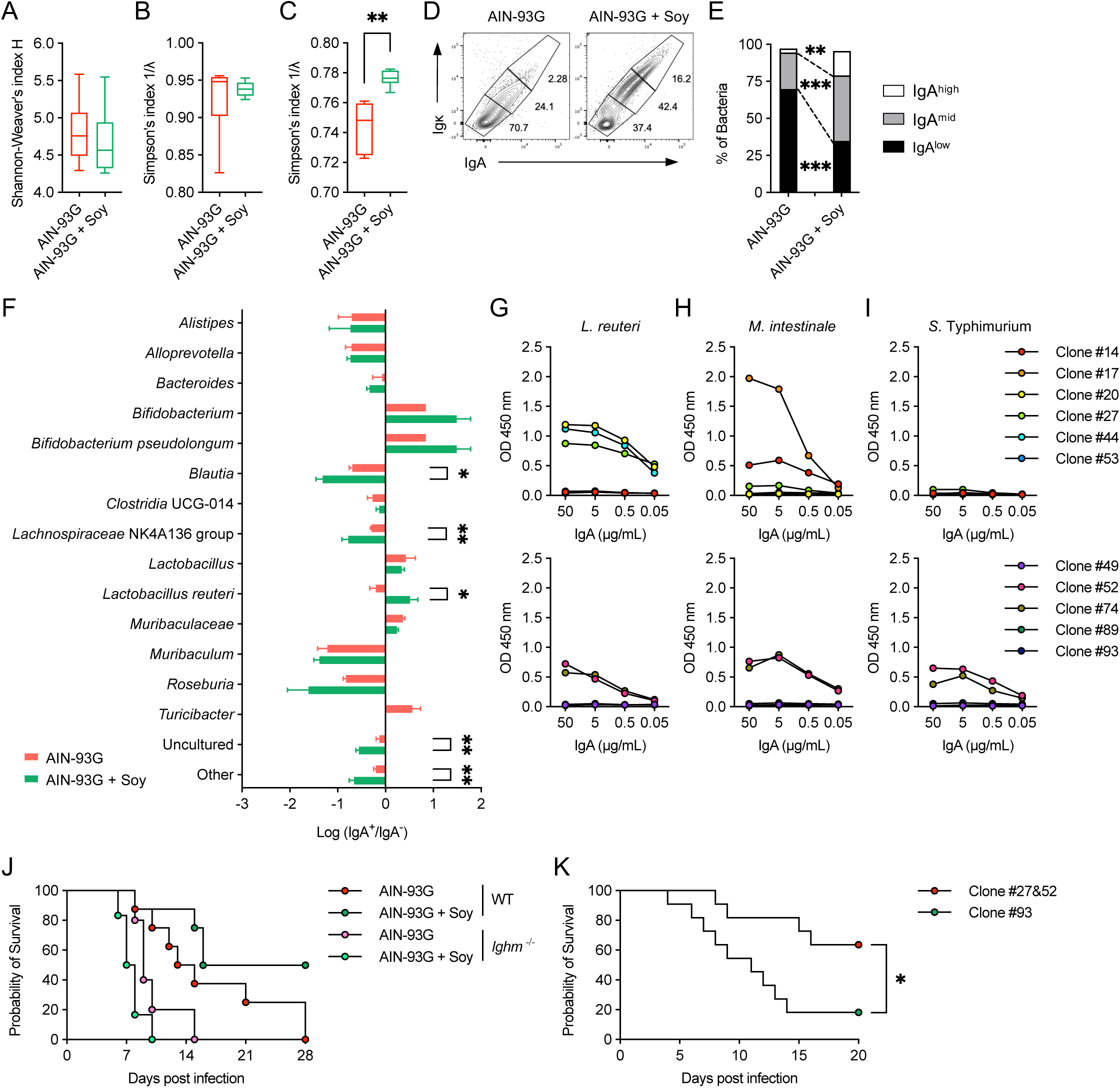
Soy-induced IgA is polyreactive and enhances resistance to *S*. Typhimurium infection. (A-C) IgH sequence analysis of IgA-producing cells from the distal small intestinal lamina propria in mice fed AIN-93G or AIN-93G + Soy diets (*n* = 6 mice/group, mean ± s.d.). (A) Diversity (Shannon-Weaver’s index). (B) Ratio of nonsynonymous substitutions (Simpson’s index). (C) Affinity maturation index (R_CDR_/S_Total_+R_CDR_). R_CDR_, replacement in CDR1-3; S_total_, silent mutations in both CDR1-3 and in FWR1-3. (D-F) Flow cytometry analysis of IgA-coated bacteria in distal small intestinal content from mice fed either the AIN-93G or the AIN-93G + Soy diet. (D and E) Representative plots of IgA staining within SYTO Red⁺ bacteria (D) and quantification of IgA^high^, IgA^mid^, and IgA^low^ bacteria (E) (*n* = 6 mice/group). (F) Log ratio of IgA-coated to IgA-uncoated bacteria (*n* = 6 mice/group). (G-I) Binding assay of culture supernatants from IgA-producing hybridomas to lysates of *L. reuteri* (G), *M. intestinale* (H), and *S*. Typhimurium (I). See also Table S4 for details of each clone. (K and L) Kaplan-Meier survival curves of mice orally infected with *S*. Typhimurium χ3181. (K) Mice were fed AIN-93G or AIN-93G + Soy diets for 2 weeks before infection (*n* = 8 mice/group). (L) *Igha*^-/-^ (IgA-deficient) mice were infected with bacteria pre-incubated with ascites containing *S*. Typhimurium-reactive IgA clones (#27 and #52) or a non-reactive IgA clone (#93) (*n* = 11-12 mice/group). Data in (A-F, J, and K) are from one representative experiment of at least two. Data in (G-I) are representative of two independent experiments. Statistical analysis was performed by (A-C, E, and F) Welch’s t-test or Student’s t-test, or (K and L) Logrank (Mantel-Cox) test. \**p* < 0.05; \*\**p* < 0.01; \*\*\**p* < 0.001. See also Figure S11 and S12.

To further characterize the reactivity of IgA produced in mice colonized with *L. reuteri* and *M. intestinale*, we generated hybridomas from intestinal lamina propria cells and obtained eleven IgA-producing clones. Sequence analysis of their IgH genes revealed that the clones possessed varying levels of somatic mutations (0.37–7.84%). Binding assays showed that five of these clones bound to *L. reuteri* and five bound to *M. intestinale*, with three clones binding to both. Three clones also recognized soy protein (Figures 6G, 6H, and S12A). Interestingly, two clones (Clone#52 and #74) exhibited binding not only to the two commensal bacteria but also to *Salmonella enterica* serovar Typhimurium (*S*. Typhimurium) (Figure 6I). This polyreactivity appeared specific, as these clones bound to keyhole limpet hemocyanin (KLH) but not to other protein antigens like flagellin or insulin (Figures S12C and S12D). Furthermore, none of the eleven clones, including those with low mutation rates, showed significant binding to TI antigens such as LPS or double-stranded DNA (dsDNA), arguing against broad, non-specific polyreactivity (Figures S12E and S12F).

Given that *S*. Typhimurium is a natural intestinal pathogen in mice, we hypothesized that the increased commensal-reactive IgA that cross-reacts with *S*. Typhimurium could offer protection against infection.^52^ Indeed, upon oral *S.* Typhimurium infection, mice fed the AIN-93G + Soy diet showed significantly prolonged survival compared to those fed the AIN-93G diet (Figure 6J). This protective effect was B cell-dependent, as it was abolished in *Ighm*^-/-^ (μMT) mice, which lack mature B cells (Figure 6J). More directly, infection with *S*. Typhimurium pre-incubated with the *S*. Typhimurium-reactive IgA clones (Clone#27 and #52) to IgA-deficient (*Igha*^-/-^) mice significantly prolonged the survival of mice compared to infection with bacteria pre-incubated with a non-reactive IgA clone (Clone#93) (Figure 6K). These findings illustrate that soy supplementation enhances the production of high-affinity, polyreactive IgA, which contributes to the host’s defense against intestinal infections by binding to and neutralizing pathogenic bacteria like *S*. Typhimurium.

## Discussion

Our findings delineate a specific diet-microbiota axis in weaning mice, where dietary soy establishes a niche for two key commensals, *L. reuteri* and *M. intestinale*, to synergistically drive mucosal IgA responses. This interaction reveals a clear division of labor: *L. reuteri* serves as a primary source of cognate antigens, while *M. intestinale* functions as a critical adjuvant by promoting IL-1β production from myeloid cells. The dominant role of *L. reuteri* as an antigen was evidenced by our finding that a significant fraction—over 60%—of the Tfh cell repertoire in soy-fed mice was reactive to this single species. This concept of keystone species driving a focused immune response aligns with the emerging paradigm that the intestinal T cell repertoire, rather than being broadly reactive, is surprisingly focused on a small number of commensal species.^53,54^ This immunological focus may represent an efficient host strategy, where T cell clonotypes recognizing broadly conserved antigens are selected to provide defense against a swath of related strains. Although the reactivity of many of our clones was of low affinity, this broad targeting of a single commensal species strongly supports the idea that *L. reuteri* provides keystone antigens in this dietary context. Our study extends this paradigm by revealing a mechanism through which such a focused immune response can be initiated: a dietary intervention that selectively expands the commensal, providing the keystone antigen. This synergistic triad—diet, antigen, and adjuvant—thus highlights a sophisticated mechanism for immune education in early life.

The necessity of this specific triad was underscored by our experiments using mice from the Inasa facility, which naturally harbor low levels of these key bacteria. In these mice, soy supplementation alone was insufficient to induce Tfh cells, demonstrating that the diet itself is not directly immunomodulatory. Conversely, oral inoculation of the two bacteria failed to promote Tfh induction without the soy diet, likely because the bacteria could not efficiently colonize the intestine. The specificity of this bacterial synergy was further highlighted by our finding that substituting *M. intestinale* with another commensal, *F. rodentium*, failed to recapitulate the Tfh-inducing effect, confirming that not just any combination of bacteria can fulfill these synergistic roles. Intriguingly, our findings from the Inasa mice also revealed that even transient exposure to the inoculated bacteria, without stable colonization, was sufficient to initiate a modest germinal center response, offering new insights into the dynamics of host-microbe interactions.

Our study also provides insight into how soy, as a dietary component, shapes this specific microbial niche. The AIN-93G diet, composed of purified ingredients, likely limits nutrient availability for gut bacteria in the distal small intestine. In contrast, soybeans, rich in prebiotics such as resistant proteins, soluble fiber, and oligosaccharides, are known to modulate the gut microbiota.^55^ Our findings align with previous research demonstrating that soy protein increases the abundance of *Muribaculaceae*,^56^ the family to which *M. intestinale* belongs. Indeed, we found that supplementation with soy protein and fiber, combined with the major soy oligosaccharides stachyose and raffinose,^48^ was required to restore the colonization of *M. intestinale*. This aligns with the known capacity of *Muribaculaceae* to hydrolyze complex carbohydrates.^47^ However, this combination was insufficient to increase *L. reuteri* abundance, suggesting that additional, yet-to-be-identified soy components are essential for supporting the growth of this keystone antigen-providing bacterium. Notably, while soy supplementation clearly rescued the phenotype, the number of IgA-producing plasma cells did not fully reach the levels observed in mice fed the unpurified CE-2 diet. This suggests that while soy is a critical driver, additional factors—whether other dietary components in the crude diet or the microbes they support—may also contribute to the robust induction of intestinal IgA. Our data reveal *M. intestinale* as a key immunogenic bacterium that functions as an adjuvant in this axis. Its ability to enhance IL-1β and IL-6 production from BMDCs is consistent with recent findings that its cardiolipin, a cell membrane component, induces pro-inflammatory cytokines from human monocytes via TLR1/2 signaling.^57^ This potent adjuvant activity may also explain the observed increase in Th17 cells in the soy-fed group. While a previous study reported an ILC2-dependent IgA induction by *M. intestinale* in the stomach,^58^ our findings reveal a distinct, PP-dependent mechanism in the small intestine where *M. intestinale* synergizes with *L. reuteri* to drive Tfh-cell differentiation and subsequent TD-IgA responses. This highlights the complex and context-dependent nature of IgA induction by this bacterium. Regarding its localization, our 16S rRNA analysis suggested a preference for the mucus-associated fraction, which may be attributed to its ability to hydrolyze host-derived mucin glycans.^47^ Such proximity to the epithelium could increase its interaction with immune cells in PPs, thereby enhancing its immunomodulatory effects. The functional consequence of this axis is the production of high-affinity, polyreactive IgA that enhances host defense. In line with the alterations in the inductive sites, we found that soy supplementation restored IgA-producing plasma cells in the lamina propria. The resulting IgA exhibited cross-reactivity against the enteric pathogen *S.* Typhimurium, and its passive administration was sufficient to protect IgA-deficient mice from lethal infection. This finding provides a critical mechanistic layer to the established importance of secretory antibodies in salmonellosis. While previous work highlighted the role of "innate" TI-IgA,^59^ our study extends this model by identifying a distinct, adaptive TD component that can be dynamically shaped by environmental factors like diet.

*L. reuteri* is a well-studied probiotic with diverse immunomodulatory functions, including the promotion of IgA production in pre-weaning mice through various mechanisms.^60^ Notably, its administration is known to boost IgA production in the small intestine specifically through the induction of Tfh and GCB cells, establishing a commensal community resilient to dysbiosis.^61^ Our study builds upon this by demonstrating that the synergy with *M. intestinale* is critical for a robust response. Furthermore, the significant IgA coating we observed on *L. reuteri* in the AIN-93G + Soy diet-fed group is also noteworthy. This observation is consistent with previous studies suggesting that IgA can function not only in pathogen elimination but also in commensal bacteria colonization. For instance, while IgA recognizing the pathobiont SFB contributes to its elimination,^62^ IgA can also act as a scaffold for other *Lactobacillus* species, assisting in their mucosal adhesion.^63^ Thus, *L. reuteri* may leverage the Tfh-IgA axis not only to train the immune system but also to secure its own niche close to the epithelium, creating a positive feedback loop for immune surveillance.

A key question is whether this process requires activation-induced deaminase (AID). While experiments in AID-deficient mice are confounded by severe dysbiosis,^18^ our finding that these protective IgA clones possess extensive somatic mutations provides the strongest possible evidence that their generation is an AID-dependent, Tfh-driven process. A fascinating aspect of our study is the polyreactivity of the IgA clones that recognize not only commensals but also the pathogen *S. Typhimurium*. The identity of the specific epitope shared among these bacteria remains an important question, but our data provide several clues. The reactivity is not directed against common microbial motifs like LPS or flagellin, and the AID-dependent nature of these IgA clones suggests recognition of a defined, likely proteinaceous, epitope. This is further supported by their binding to the protein antigen KLH. Pinpointing the precise molecular targets of these protective, polyreactive IgA clones is a critical direction for future research and will be key to harnessing this phenomenon for therapeutic or vaccine strategies.

The induction of this axis appears to be initiated by M-cell-mediated uptake of luminal components, as the Tfh response was diminished in M-cell-deficient mice. The subsequent expansion of GCB and IgA⁺ B cells was entirely dependent on the initial Tfh cell differentiation, as this effect was completely abrogated in Tfh-deficient mice, confirming the specificity of the Tfh-IgA axis rather than a general increase in PP cellularity.

It is important to note that our study was conducted during the "weaning reaction," a critical window of immunological plasticity.^43^ Our findings clearly demonstrate that the diet-microbiota-Tfh axis is highly plastic and responsive during this formative period. This responsiveness, however, also highlights its dependence on continuous stimulation; cessation of soy intake led to a rapid decline in Tfh and IgA⁺ B cells to baseline levels. This suggests that sustained dietary input is required to maintain the microbial community and the associated immune response. A key remaining question is whether similar plasticity exists in mature adults, whose immune systems may be more resilient to dietary shifts.

While our study was conducted entirely in mice, its findings have potential human relevance. The key components of this axis—soy, *L. reuteri*, and bacteria from the Muribaculaceae family—are all relevant to human diets and microbiota.^47,64^ Although *M. intestinale* is considered mouse-specific, its immunomodulatory lipid activates human cells, suggesting the crucial factor is the function of providing a TLR1/2-ligand adjuvant, not the specific bacterial species.^57^ It is therefore plausible that a "functional equivalent" to *M. intestinale* could complete a similar synergistic triad in the human gut. In conclusion, our study underscores the potential of dietary interventions and probiotic strategies in modulating the gut microbiota and maturing the mucosal immune system.

## Acknowledgment

We thank the Animal Facility at the Keio University Faculty of Pharmacy for breeding and maintaining the mice, and the Instrument Management Core at the Keio University Faculty of Pharmacy for their support. We thank Ryutaro Tamano for technical assistance and Peter Burrows for editing this manuscript.

## Author contributions

D.T. and K.H-M. conceived the study and designed the experiments. D.T. primarily wrote the manuscript. D.T. and K.H-M. prepared all figures and tables with assistance from H.M.. K.H-M. performed most of the pre-revision experiments, while H.M. and D.T. performed most of the experiments for the revision. D.T., K.H-M., and H.M. were the primary individuals responsible for data analysis and interpretation, with assistance from Y.Ki., K.H., Y.Ka., Y.F., S.T., H.O., Y.O., Y. KO., E.H., K.T., Se.K., N.M., H.Y-A., Y.O., Y.S., W.O., Sh.K.. D.T. performed the single-cell RNA-seq analysis. K.H-M., Y.Ki., K.S., and N.M. maintained the germ-free mice and conducted infection experiments. D.T. and K.H. supervised the project. All authors reviewed and approved the final manuscript.

## STAR Methods

### Contact for reagent and resource sharing

Further information and requests for reagents may be directed to, and will be fulfilled by, Daisuke Takahashi, and Koji Hase

### Mice and experimental treatments

All animal experiments were approved by the Ethics Committee of Keio University (#A2022-283) and conducted in accordance with institutional guidelines. Male mice were used for all experiments. Mice were housed at the Animal Care Facility of the Faculty of Pharmacy, Keio University, under specific pathogen-free (SPF) conditions with free access to water and irradiated feed (30 kGy gamma radiation). Germ-free (GF) mice were maintained in sterile vinyl isolators with free access to sterile water and autoclaved feed. All mice were housed under controlled conditions (21–22°C, 12-hour light/dark cycle). C57BL/6JJcl mice (from the Ishibe Breeding Farm unless otherwise specified) and BALB/cAJcl-nu/nu mice were purchased from CLEA Japan (Tokyo, Japan). C57BL/6JJmsSlc mice (from the Inasa Breeding Farm) were purchased from Japan SLC, Inc. (Shizuoka, Japan). GF ICR mice were purchased from Sankyo Labo Service Corporation (Tokyo, Japan). *Bcl6*-tdTomato-creERT2 mice were a kind gift from Dr. Yosuke Harada (Tokyo University of Science).^65^ *Myd88*^⁻/⁻^*Ticam1*^⁻/⁻^ mice were purchased from Oriental BioService, Inc. (Kyoto, Japan). *Cd4*-cre, *Bcl6*^fl/f^, *Vil1*-cre, and *Tnfrsf11a* ^fl/f^ transgenic mice were purchased from The Jackson Laboratory (Bar Harbor, ME, USA). The experimental diets included the unpurified diet CE-2 (CLEA Japan) and the semi-purified diet AIN-93G (Oriental Yeast, Tokyo, Japan). Modified AIN-93G diets were supplemented with soy powder (Pioneer Confectionery & Baking, Kanagawa, Japan), soy protein (Fuji-pro F; Fuji Oil, Osaka, Japan), soy fiber (Soyafibe-S; Fuji Oil), stachyose hydrate (BIOSYNTH Carbosynth, Compton, UK), or D-(+)-Raffinose Pentahydrate (Nacalai Tesque, Kyoto, Japan). The detailed compositions of all diets are provided in Tables S1, S2, and S3. Unless otherwise specified, experimental diets were provided to mice starting at 3 weeks of age, while GF mice were used for experiments at 4–5 weeks of age. For antibiotic administration, mice were given drinking water containing vancomycin (VCM; 0.5 g/L; FUJIFILM Wako Pure Chemical Corporation, Osaka, Japan), streptomycin (SM; 1 g/L; Nacalai Tesque), or erythromycin (EM; 1 g/L; Nacalai Tesque). At the end of the experiments, mice were euthanized by cervical dislocation under isoflurane anesthesia, and tissues such as the small intestine and Peyer’s patches were immediately collected for analysis.

### Preparation of mouse bone-marrow-derived DCs (BMDCs)

The limbs were harvested from male mice aged 6 to 9 weeks. The bones were immersed in a 70% ethanol solution after the skin and muscle were removed. Subsequently, bone marrow (BM) cells were extracted from the bones and suspended in phosphate-buffered saline (PBS; pH 7.2, Nacalai tesque) containing 2% fetal bovine serum (FBS; BioWest, Riverside, MO) and 2mM EDTA (Nacalai Tesque). The collected cell suspension, with a count of 1 × 10^7^ cells per dish, was cultured in RPMI-1640 medium (PBS; pH 7.2, Nacalai Tesque) containing 10% FBS (BioWest), GlutaMAX (Thermo Fisher Scientific, Waltham, MA), Penicillin-Streptomycin Mixed Solution (Nacalai Tesque), 55 µM mercaptoethanol, and 12.5 mM HEPES (pH 7.2, Nacalai Tesque). The culture medium was further supplemented with 10 ng/ml recombinant mouse granulocyte-macrophage colony-stimulating factor (GM-CSF) (Biolegend in San Diego, CA), 5 ng/ml FMS-like tyrosine kinase 3 ligand (FLT3L) (Biolegend), and 10 ng/ml interleukin 4 (IL-4) (Biolegend). This culture period spanned 6 days, following the protocol outlined in Isobe et al. (2020).^66^ The medium was refreshed on days 3 and 5. After 6 days of culture, the cells were stimulated with 100 ng/ml lipopolysaccharides (LPS) for 24 hours to conduct the Tfh hybridoma assay.

### Flow cytometry and cell sorting

Peyer’s patches of the proximal and distal half of the small intestine were harvested macroscopically. Single leukocyte suspensions of the Peyer’s patches were prepared by mechanically disrupting tissues through 100-µm nylon mesh cell strainers (Greiner Bio-One, Kremsmünster, Austria) in 2% FCS RPMI1640 media (Nacalai Tesque, Kyoto, Japan). Leukocytes were pre-incubated for 15 minutes on ice with a monoclonal antibody (mAb) against CD16/32 (S17011E; BioLegend, San Diego, CA, USA) in 2% FCS and 0.1% NaN3 in PBS (Nacalai Tesque) before surface antigen staining. Cell staining was performed with mAbs including Brilliant Violet 786 (BV786)-conjugated anti-CD4 (RM4-5; BD Biosciences, Franklin Lakes, NJ, USA), BV605-conjugated anti-TCRβ-chain (H57-597; BD Biosciences), BV650-conjugated anti-CD25 (PC61; BD Biosciences), BV 510-conjugated anti-CD45 (30-F11; BioLegend), Pacific Blue-conjugated anti-GL-7 (GL7; BioLegend), PerCP-eFluor 710-conjugated anti-PD-1 (J43; Thermo Fisher Scientific, Waltham, MA, USA), FITC-conjugated anti-IgA (C10-3; BD Biosciences), APC-R700-conjugated anti-CD19 (1D3; BD Biosciences), APC-conjugated anti-CXCR5 (L138D7; BioLegend), and followed by dead cell staining with Fixable Viability Stain 780 (FVS780; BD Biosciences). The cells were then fixed, permeabilized, and stained with mAbs, including PE-conjugated anti-Foxp3 (R16-715; BD Biosciences) and PE-CF594-conjugated anti-Bcl-6 (K112-91; BD Biosciences) using a transcription factor buffer set (BD Biosciences). Flow cytometry was performed using a FACSCelesta flow instrument with DIVA v9.0 (BD Biosciences), and cell sorting was performed using a FACSAria III flow instrument, and data were analyzed using FlowJo version 10.9 (BD Biosciences).

### T cell hybridoma generation and screening

FACS-purified PP TCRβ^+^CD4^+^hCD2^-^CD25^-^Bcl-6-tdTomato^+^ T cells isolated from PPs of AIN-93G + Soy diet-fed Bcl6-tdTomato-cre^ERT2^ mice were stimulated *in vitro* for 3 days in tissue culture plates coated with 5 µg/ml each of anti-CD3 and anti-CD28 mAbs, fused with Nur77-Egfp^+^αβTCR^-^ BW5147 thymoma cells by using GenomONE-CF (ISHIHARA SANGYO KAISHA, LTD., Osaka, Japan) and selected in Hypoxanthine-aminopterin-thymidine (HAT; Sigma-Aldrich, St. Louis, MO, USA) containing complete RPMI1640 medium. The response of a mixture of hybridoma clones toward autoclaved bacterial lysates was assessed by the concentration of IL-2 in the supernatant. In brief, 2×10^4^ hybridoma cells were incubated with 2×10^3^ bone-marrow-derived dendritic cells alone (no antigen control) or with small intestinal contents from GF, *L. retuteri*, or *M. intestinale* gnotobiotic mice. After 24 hr, the amount of secreted IL-2 was measured by ELISA MAX Deluxe Set (BioLegend) following the manufacturer’s instructions.

### Bacterial culture and generation of gnotobiotic mice

The following bacterial strains were used: *Muribaculum intestinale* (strains YL5, YL7, YL27), *Limosilactobacillus reuteri* (an in-house strain), and *Faecalibaculum rodentium* (JCM 30274). For cultivation, *M. intestinale* was plated on Gif anaerobic medium (GAM; Nissui, Tokyo, Japan) agar supplemented with 1% stachyose (Biosynth, Berkshire, UK) and incubated for 3 days at 37°C under anaerobic conditions. *L. reuteri* was plated on De Man-Rogosa-Sharpe (MRS; BD Biosciences, Franklin Lakes, NJ, USA) agar and incubated for 2 days at 37°C anaerobically or aerobically. *F. rodentium* was plated on GAM agar and incubated for 2 days at 37°C anaerobically. For liquid culture, single colonies were picked and suspended in their respective broths (GAM + 1% stachyose, MRS, or GAM) and incubated for 24–48 hours at 37°C. For *F. rodentium*, a pre-culture was grown for 2 days, followed by a main culture for 1 day under the same conditions. For cryopreservation, bacterial cultures were centrifuged at 10,000 × g for 5 min at 4°C, and the cell pellets were resuspended in their respective broths containing 15–30% glycerol and stored at −80°C. To generate gnotobiotic mice, germ-free (GF) mice were colonized by oral gavage with specific bacterial suspensions. The colonization protocols varied depending on the experiment: For the experiments shown in Figure 4A–G, GF mice fed an AIN-93G + Soy diet were colonized with *M. intestinale* (1 × 10¹⁰ CFU, administered three times over 3 days), *L. reuteri* (1 × 10⁹ CFU, administered once), or both. For the experiments shown in Figure S9, GF mice fed an AIN-93G + Soy diet were co-colonized with either *L. reuteri* and *F. rodentium* (1 × 10⁸ CFU each) or *L. reuteri* and *M. intestinale* (1 × 10⁸ CFU each). The bacterial mixtures were administered six times over 6 days.

### Stimulation of BMDCs with bacterial lysates and measurement of cytokine levels

Cultured *M. intestinale* and *L. reuteri* were autoclaved at 121°C for 20 minutes and then centrifuged to remove debris, preparing bacterial lysates. Bone marrow-derived dendritic cells plated at 1×10^5^ cells/well were treated with bacterial lysates at a final protein concentration of 5 µg/ml, and LPS at a final concentration of 100 ng/mL. After 24 hours, culture supernatants were collected and IL-6 levels in the supernatants were measured using the ELISA MAX Standard Set (BioLegend) according to the manufacturer’s protocol. For IL-1β measurement, 1 µM nigericin (InvivoGen, San Diego, USA) was added 10 hours after stimulation with LPS and bacterial lysates, and culture supernatants were collected after 14 hours. The ELISA MAX Standard Set (BioLegend) was used to measure IL-1β and IL-6 levels in culture supernatants according to the manufacturer’s protocol.

### Quantification of IgA by Enzyme-Linked Immunosorbent Assay (ELISA)

Intestinal contents and feces were collected from 7-week-old mice and suspended in PBS containing Complete Proteinase Inhibitor Cocktail (Roche, Mannheim, Germany). The suspension was centrifuged at 12,000×g for 5 minutes at 4°C. The supernatant was diluted with 2% bovine serum albumin (BSA; globulin-free, Nacalai Tesque) in PBS. Flat-bottom 96-well MaxiSorp Nunc-Immuno plates (Thermo Fisher Scientific) were coated with goat anti-mouse IgA antibody (Bethyl Laboratories, TX, USA) at room temperature for 1 hour, then washed 4 times with 0.1% Tween 20 Tris-buffered saline (TBS-T). The plates were blocked with 2% BSA/PBS at room temperature (20-25°C) for 1 hour. After washing the plates 4 times with TBS-T, HRP-conjugated goat anti-mouse IgA antibody (Bethyl Laboratories) was incubated at room temperature for 1 hour. The plates were washed 5 times with TBS-T, followed by incubation with 1-Step Ultra TMB-ELISA Substrate Solution (Thermo Fisher Scientific) at room temperature for up to 15 minutes, and the reaction was stopped by adding 1.2M sulfuric acid. Absorbance was measured at 450 nm using an Infinite 200 PRO microplate reader (Tecan, Männedorf, Switzerland).

### IgA-bacterial flow cytometry

Mouse feces were crushed on a 70-μm cell strainer (Greiner). After centrifugation at 500 × g for 5 minutes at 4°C to remove large debris, the supernatant was centrifuged at 12,000 × g for 5 minutes to pellet the bacteria. The bacterial pellet was resuspended in 1% BSA/PBS containing biotinylated goat anti-mouse IgA (BD Biosciences). After washing with 1% BSA/PBS, the bacteria were stained with streptavidin-BV421 (BioLegend). Subsequently, the bacteria were washed by centrifugation at 12,000 × g for 5 minutes and resuspended in 1% BSA/PBS containing 50 nM SYTO Red (Thermo Fisher Scientific). Flow cytometry analysis was performed using a FACSCelesta flow cytometer equipped with DIVA v9.0 (BD Biosciences) and data were analyzed using FlowJo version 10.9 (BD Biosciences). IgA-binding bacteria were magnetically isolated using the MojoSort system (BioLegend) with streptavidin nanobeads (BioLegend) according to the manufacturer’s protocol. To isolate IgA-coated bacteria, the bacterial pellet, after staining with biotinylated goat anti-mouse IgA (BD Biosciences), was incubated with MojoSort Streptavidin Nanobeads (BioLegend). IgA-positive bacteria bound to nanobeads were separated using a MojoSort magnet (BioLegend) according to the manufacturer’s instructions. The bead-bound fraction containing IgA-coated bacteria was collected, washed twice with 1% BSA/PBS, and processed for DNA extraction and 16S rRNA gene sequencing.

### DNA extraction and microbiome genome sequencing

Genomic DNA was extracted from small intestinal contents or mucus samples with a QIAamp Fast DNA Stool Mini Kit (Qiagen, Hilden, Germany) for Figure 1C and Figure 2I, or a QIAamp PowerFecal Pro DNA Kit (Qiagen) for all other analyses in accordance with the manufacturer’s protocols. A 16S rRNA genomic libraries were constructed in accordance with the protocol of the Illumina technical note with some modifications. Briefly, the extracted DNA samples were used as DNA templates in a polymerase chain reaction (PCR) carried out in a reaction mixture of template DNA, KAPA HiFi HotStart Ready Mix (Roche), and primers specific for the 16S rRNA V3–V4 region under the following conditions: initial denaturation at 95 °C for 3 min, followed by 35 cycles at 95 °C for 30 s, 55 °C for 30 s, and 72 °C for 30 s. A final elongation step was performed at 72 °C for 5 min. The amplicons were purified with AMPure XP beads (Beckman Coulter, Brea, CA, USA) and attached to dual indices by index PCR with a Nextera XT Index kit (Illumina, San Diego, CA, USA). The libraries were purified with AMPure XP beads (Beckman Coulter). The purified libraries were diluted to 4 nM with Tris–HCl buffer and pooled. The libraries were then sequenced on a Miseq (Illumina) with 300 bp paired-end reads. Shotgun metagenome sequencing libraries were prepared using the NEBNext Ultra II FS DNA Library Prep Kit for Illumina (New England Biolabs, MA, USA) according to the manufacturer’s protocol and dual-indexed with NEBNext Multiplex Oligos for Illumina (New England Biolabs, MA, USA), then purified with AMPure XP beads. The libraries were pooled and sequenced with 150 bp paired-end reads on a HiSeq platform (Illumina). Shotgun metagenome sequencing libraries were prepared using the NEBNext Ultra II FS DNA Library Prep Kit for Illumina (New England Biolabs, MA, USA) according to the manufacturer’s protocol and dual-indexed with NEBNext Multiplex Oligos for Illumina (New England Biolabs, MA, USA), then purified with AMPure XP beads. The libraries were pooled and sequenced with 150 bp paired-end reads on a HiSeq platform (Illumina).

### Quantification of bacteria by qPCR

Bacterial genomic DNA was isolated from fecal pellets and intestinal luminal contents using a QIAamp PowerFecal Pro DNA Kit (Qiagen, Hilden, Germany). Real-time quantitative PCR (qPCR) was used to quantify the genomic DNA of specific bacterial taxa (SFB, *L. reuteri*, *M. intestinale*, *F. rodentium*) and the total bacterial load. Probe-based quantification of SFB, *L. reuteri*, and *M. intestinale* was performed using taxon-specific primers and probes (Supplementary Table 5). Each 20 µL reaction contained 2 µL of template DNA (5 ng/µL), 10 µL of 2x Buffer for rTth/TTx (DNA), 0.25 µL of Hot Start TTx DNA Polymerase (Toyobo, Osaka, Japan), 0.5 µL of the probe, 0.5 µL each of forward and reverse primers, and 6.25 µL of sterile water. The qPCR was performed on a CFX real-time PCR system using CFX Maestro Software (Bio-Rad, Hercules, CA, USA) with the following program: an initial denaturation at 95°C for 3 min, followed by 50 cycles of 95°C for 15 s, 50°C for 20 s, and 72°C for 10 s. SYBR Green-based quantification of *F. rodentium* was performed using specific primers (Supplementary Table 5). Each 20 µL reaction consisted of 2 µL of template DNA (5 ng/µL), 10 µL of KOD SYBR qPCR Mix (Toyobo), 0.4 µL each of forward and reverse primers, and 7.2 µL of sterile water. The qPCR was performed on a CFX Connect real-time PCR analysis system (Bio-Rad) with the following program: 98°C for 3 min, followed by 40 cycles of 98°C for 10 s, 52°C for 10 s, and 68°C for 30 s. Absolute quantification for each bacterium was accomplished using standard curves. For SFB, a standard was prepared by cloning a PCR-amplified fragment (primers in Supplementary Table 5) into a pCR-Blunt vector (Thermo Fisher Scientific, Waltham, MA, USA). For *L. reuteri*, *M. intestinale*, and *F. rodentium*, standards were prepared by extracting genomic DNA from a known number of colony-forming units (CFU) of *in vitro* cultured bacteria. Quantification of total bacterial load was performed using a SYBR Green-based assay with universal 16S rRNA primers (5ʹ-CCTACGGGNGGCWGCAG-3ʹ and 5ʹ-GACTACHVGGGTATCTAATCC-3ʹ), as previously described.^67^ The qPCR was performed on a CFX Connect system (Bio-Rad) with the following program: 98°C for 3 min, followed by 40 cycles of 94°C for 15 s, 55°C for 10 s, and 60°C for 1 min, and completed with a melting curve analysis. A standard curve was generated using a dilution series (10¹ to 10⁸ CFU) of *E. coli* DNA.

### Microbiome analysis

For 16S rRNA gene sequencing, after removing mouse-associated contaminants using Bowtie2^68^ with the mm10 index, FASTQ files were analyzed using the QIIME2 pipeline (QIIME2 version 2020.2).^69^ After conversion to the qza format, the sequence data were demultiplexed and summarized using QIIME2 paired-end-demux. Then, the sequences were trimmed and denoised with the DADA2^70^ plugin for QIIME2. Taxonomic assignment was performed using a naïve Bayes fitted classifier trained on the SILVA database (version 138)^71^ with the feature-classifier plugin for QIIME2. Diversity analysis was performed with QIIME2 coremetrics-phylogenetic. The relative abundances of each taxon were calculated using the taxa collapse QIIME2 plugin. For Shotgun metagenome sequencing, analysis was performed using tools included in the bioBakery suite (https://github.com/biobakery). Quality filtering to remove low-quality bases and adapter sequences, as well as mapping to the host genome sequences, was conducted using Kneaddata, and bacterial community composition analysis was performed using MetaPhlAn. ^72,73^

### Generation of IgA-producing hybridomas

B cell hybridomas were generated from the small intestinal lamina propria as previously described, with some modifications.^14^ Briefly, small intestines were collected from GF ICR mice that had been fed an AIN-93G + Soy diet and colonized with *L. reuteri* and *M. intestinale*. The intestines were opened longitudinally, washed with PBS to remove luminal contents, and then shaken in PBS containing 5 mM EDTA for 20 min at 37°C to remove epithelial cells. The remaining lamina propria tissue was minced and digested in RPMI-1640 medium containing 2% fetal bovine serum, 38.5 µg/mL Liberas TM (Roche, Basel, Switzerland), and 125 µg/mL DNase I (Sigma-Aldrich, St. Louis, MO, USA) for 1 h at 37°C with shaking.

The isolated lamina propria cells were washed with PBS and fused with NS-1 myeloma cells using polyethylene glycol. Cell fusion and subsequent subcloning were performed according to the manufacturer’s protocol (ClonaCell-HY Hybridoma Cloning Kit; STEMCELL Technologies, Vancouver, Canada). IgA-secreting hybridoma clones were identified and selected by a standard sandwich ELISA using an IgA ELISA kit (Bethyl Laboratories, Montgomery, TX, USA). IgA-producing hybridomas were selected using IgA ELISA

### Sequencing and analysis of immunoglobulin heavy chains (IgH)

The sample processing and analysis were performed as previously described with some modifications.^74^ Briefly, RNA was extracted from a 5-mm segment of the distal small intestinal tissue using NucleoSpin RNA PLUS (Takara Bio, Shiga, Japan). Subsequently, IgA cDNA was synthesized using SuperScript IV Reverse Transcriptase and the following mix of three gene-specific primers (5’-ATCAGGCAGCCGATTATCAC-3, 5’-TCTCCTTCTGGGCACTCG-3’, and 5’-TGAATGATGCGCCACTGT-3’). To generate template libraries, one forward primer and seven reverse primers for rearranged IgA sequences^19^ were modified to incorporate adaptor sequences for Illumina indexes (where ‘S’ is C or G; ‘R’ is A or G; ‘N’ is A, G, C or T; ‘M’ is A or C; and ‘W’ is A or T): IgA, Fw, 5’-TCGTCGGCAGCGTCAGATGTGTATAAGAGACAGGAGCTCGTGGGAGTGTCAGTG-3’, Rv1, GTCTCGTGGGCTCGGAGATGTGTATAAGAGACAGGASARGTNMAGCTGSAGSAGTC, Rv2, GTCTCGTGGGCTCGGAGATGTGTATAAGAGACAGGASARGTNMAGCTGSAGSAGTCW, Rv3, GTCTCGTGGGCTCGGAGATGTGTATAAGAGACAGGACAGGTTACTCTGAAAGWGTSTG, Rv4, GTCTCGTGGGCTCGGAGATGTGTATAAGAGACAGGAGAGGTCCARCTGCAACARTC, Rv5, GTCTCGTGGGCTCGGAGATGTGTATAAGAGACAGGACAGGTCCAACTVCAGCARCC, Rv6, GTCTCGTGGGCTCGGAGATGTGTATAAGAGACAGGAGAGGTGAASSTGGTGGAATC, Rv7, GTCTCGTGGGCTCGGAGATGTGTATAAGAGACAGGAGATGTGAACTTGGAAGTGTC). The PCR program consisted of an initial denaturation at 95°C for 4 min, followed by 34 cycles of 94°C for 30 s, 62°C for 30 s, and 72°C for 35 s, with a final extension at 72°C for 5 min. Amplicon purification, index PCR, and sequencing were performed using the same methods as for 16S rRNA sequencing. For IgH sequencing from hybridoma cells, IgA cDNA was amplified using KAPA HiFi HotStart Ready Mix (Roche) and sequenced by conventional Sanger sequencing. The resulting sequences were analyzed using the ASAP (A web server for Immunoglobulin-Seq Analysis Pipeline^75^ and MiXCR software.^76,77^

### IgA Binding assay to bacteria and antigens

For bacterial binding assays, *M. intestinale*, *L. reuteri*, and *S. Typhimurium* were cultured for 24-48 hours at 37°C in appropriate media under anaerobic or aerobic conditions. Bacteria were harvested by centrifugation, washed with PBS, and resuspended in 0.05 M Na₂CO₃ buffer. 96-well Flat-bottomed MaxiSorp Nunc-Immuno plates (Thermo Fisher Scientific) were coated with the bacterial suspension (1×10⁸ cells/well, except for *S. Typhimurium* at ∼1×10⁹ cells/well). For soluble antigen binding assays, plates were coated with various antigens diluted in 0.05 M Na₂CO₃ buffer. The following antigens were used: Soybean flour, Type I (Sigma-Aldrich, St. Louis, MO, USA) at 1 ng/well; keyhole limpet hemocyanin (KLH; Sigma-Aldrich) at 10 µg/mL; flagellin from *S. Typhimurium* (InvivoGen, San Diego, CA, USA) at 2 µg/mL; insulin (Eli Lilly, Indianapolis, IN, USA) at 0.1 unit/well; LPS from *E. coli* (InvivoGen) at 10 µg/mL; and calf thymus DNA (Rockland Inc., Limerick, PA, USA) at 10 µg/mL. All plates were incubated overnight at 4°C. Following overnight coating, plates were washed four times with Tris-buffered saline containing 0.1% Tween 20 (TBS-T) and blocked with 150 µL/well of 2% PVDF blocking buffer (Toyobo) for 1 hour at room temperature. After washing four times with TBS-T, 100 µL/well of hybridoma culture supernatants (IgA clones), diluted in Solution 1 (Can Get Signal Immunoreaction Enhancer Solution, Toyobo), were added to the antigen-coated wells. Samples were typically assayed starting at 5 µg/mL, followed by three additional 1:5 serial dilutions. Plates were incubated for 1 hour at room temperature. After incubation with the primary antibody, plates were washed four times with TBS-T. Then, 100 µL/well of HRP-conjugated goat anti-mouse IgA antibody (Bethyl Laboratories, Montgomery, TX, USA), diluted in Solution 2 (Toyobo), was added and incubated for 1 hour at room temperature. The plates were then washed five times with TBS-T, and the signal was developed by adding 1-Step Ultra TMB-ELISA Substrate Solution (Thermo Fisher Scientific). The reaction was allowed to proceed in the dark at room temperature for up to 5 minutes and was stopped by adding 1.2 M sulfuric acid. Absorbance was measured at 450 nm using an Infinite 200 PRO microplate reader (Tecan, Männedorf, Switzerland). Purified Mouse IgA κ Isotype Control antibody (BD Pharmingen, San Jose, CA, USA) was used to generate a standard curve on each plate for inter-plate normalization.

### Production and purification of monoclonal IgA from ascites

Seven-week-old female BALB/c nude mice (BALB/cAJcl-nu/nu) were intraperitoneally (i.p.) injected with 0.5 mL of pristane (Sigma-Aldrich, St. Louis, MO, USA). Two to three weeks later, the mice were i.p. injected with 1 × 10⁷ IgA-producing hybridoma cells, which had been washed and resuspended in 0.2 mL of sterile HBSS (-). Ascites fluid was collected 2–4 weeks after cell injection. The fluid was centrifuged at 1,200 rpm for 10 min at 4°C, and the supernatant was passed through a 0.45 µm filter before being stored at −80°C. Frozen ascites fluid was thawed at room temperature, clarified again by centrifugation and filtration (0.45 µm), and then diluted 10-fold with sterile PBS. To deplete IgG, the diluted ascites was applied to a HiTrap Protein G HP column (Cytiva, Marlborough, MA, USA) equilibrated with binding buffer (20 mM sodium phosphate, pH 7.0) at a flow rate of 1 mL/min. The flow-through fraction, containing the IgA, was collected. The IgA-containing fraction was then concentrated and buffer-exchanged into PBS using an Amicon Ultra-15 centrifugal filter unit with a 30 kDa molecular weight cutoff (Ultracel-30; Millipore, Burlington, MA, USA). The concentrations of IgA and any residual IgG in the purified samples were quantified by ELISA. Purified antibody preparations with confirmed IgG depletion (from clones #27, #52, and #93) were used for the *in vivo* infection experiments.

### *In vivo Salmonella* infection experiments

Three-week-old male C57BL/6J mice were fed either an AIN-93G or AIN-93G + Soy diet for three weeks. Mice were then orally infected with 1 × 10⁶ CFU of *S.* Typhimurium strain χ3181, and their survival was monitored daily. For the B cell-deficient control group, B cell-deficient (*Ighm*⁻/⁻; µMT) mice were used. To deplete their commensal microbiota, µMT mice were orally administered an antibiotic cocktail containing ampicillin (1 mg/mL; Nacalai Tesque, Kyoto, Japan), neomycin (1 mg/mL; Nacalai Tesque), vancomycin (0.5 mg/mL; FUJIFILM Wako Pure Chemical Corporation, Osaka, Japan), and metronidazole (0.5 mg/mL; FUJIFILM Wako Pure Chemical Corporation). Following antibiotic treatment, these mice were infected with *S. Typhimurium* as described above. To directly assess the protective capacity of the monoclonal IgA, an *in vivo* passive immunization experiment was performed. Purified monoclonal IgA from high-binding clones (#27 and #52) was combined (45 µg each, for a total of 90 µg/mouse). As a control, IgA from a low-binding clone (#93) was used at 90 µg/mouse. The antibody preparations were incubated with 1 × 10⁶ CFU of *S. Typhimurium* χ3181 for 1 hour at room temperature. The IgA-bacteria mixtures were then orally administered to IgA-deficient (*Igha*⁻/⁻) mice, which had been maintained on an AIN-93G + Soy diet from 4 to 8 weeks of age. Survival was monitored daily.

Single-cell RNA sequencing

Single-cell suspensions of PPs were prepared as described above. After surface staining CD11c, live CD11c^+^ cells were sorted by a FACSAria III instrument (BD Biosciences). FACS-purified CD11c^+^ cells were subjected to the preparation of 3’ Gene Expression libraries using a Chromium Next GEM Single Cell 3’ Reagent Kits v2 (Dual Index) (10X Genomics, Pleasanton, CA) following the manufacturer’s instructions. Pooled libraries were sequenced on an Illumina NovaSeq 6000 (150 bp×2 paired-end), aiming at 800.0 M reads per sample. The sequenced paired reads were processed with the “cellranger multi” function using CellRanger v8.0.0 (10X Genomics) for alignment, filtering, barcode counting, and UMI counting. Seurat objects were generated from the filtered CellRanger output using the “Read10X” function and the “CreateSeuratObject” function (Seurat v5.1.0).^78^ Low-quality cells were excluded based on the following criteria: number of genes expressed < 500, number of genes expressed > 3500, > 5% mitochondrial RNA in the total UMI counts. The Seurat objects were integrated following the SCTransform integration workflow.^79^ The number of features in the “SelectIntegrationFeatures” function was set to infinity to obtain as many variable genes as possible. Otherwise, the default parameter was set.

### Statistical analysis

Statistical analyses were performed using GraphPad Prism 10 software (GraphPad Software, La Jolla, CA, USA). A P-value < 0.05 was considered statistically significant. The specific statistical test used for each experiment is detailed in the corresponding figure legend. In general, for comparisons between two groups, a two-tailed Student’s t-test was used for data assumed to have equal variances, while Welch’s t-test was applied for data with unequal variances. For non-normally distributed data, the non-parametric Mann-Whitney U test was used. For comparisons among three or more groups, one-way or two-way analysis of variance (ANOVA) was performed. Post-hoc multiple comparisons were conducted using Tukey’s test (for all pairwise comparisons), Dunnett’s test (for comparisons against a single control group), or Šídák’s test, as appropriate. Survival curves were generated using the Kaplan-Meier method and compared using the Log-rank (Mantel-Cox) test. Data are typically presented as mean ± s.d. The experiments were not randomized, and the investigators were not blinded to allocation during experiments or outcome assessment.

