## Supplemental FIgures and Tables for "Dietary soy shapes murine microbiota to consolidate the mucosal IgA response through T follicular helper cells"

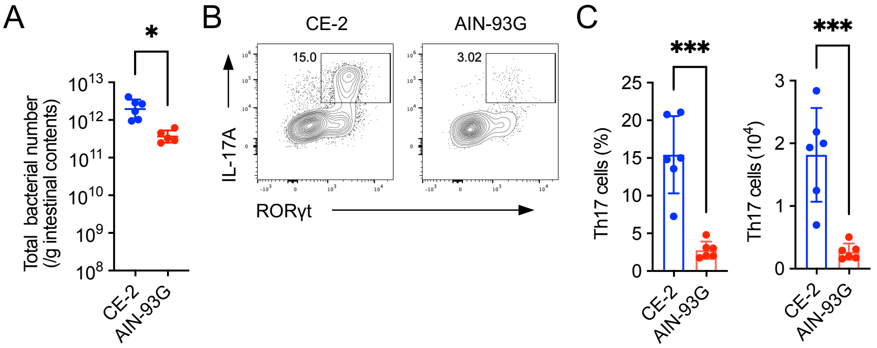


**Figure S1. A semi-purified diet reduces the total bacterial number and LP Th17 cells**

(A) Total bacterial number in the distal small intestinal content of mice fed either the CE-2 or the AIN-93G diet for 4 weeks starting from 3 weeks of age, assessed by qPCR for the 16S rRNA gene ($n$ = 6 mice/group).

(B and C) IL-17A$^+$RORγt$^+$ Th17 cells in the distal small intestinal lamina propria of mice fed either the CE-2 or the AIN-93G diet. Representative flow cytometry plots of IL-17A and RORγt staining within the CD4$^+$ T cell gate are shown in (B), and quantification of frequency and number of Th17 cells is shown in (C) ($n$ = 6 mice/group, mean ± s.d.).

Data are representative of at least two independent experiments. Statistical analysis was performed by (A, C) Welch's t-test. *$p < 0.05$; ***$p < 0.001$.

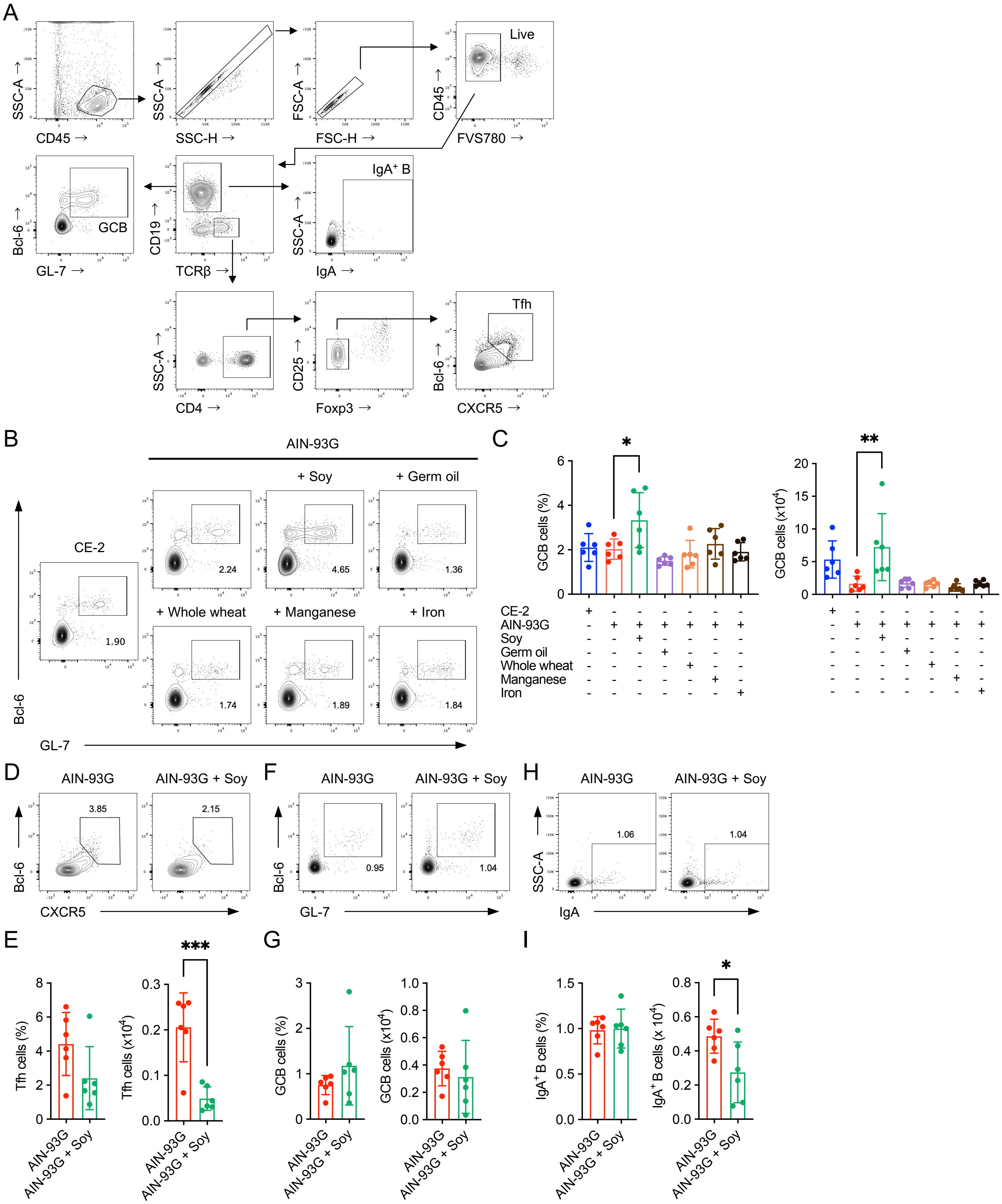


**Figure S2. Soy supplementation induces GCB, Tfh, and IgA$^+$ B cells in a commensal microbiota-dependent manner**

(A) Representative flow cytometry gating strategy for the identification of Tfh, GCB, and IgA$^+$ B cells in Peyer's patches.

(B and C) GL-7$^+$Bcl-6$^+$ GCB cells in distal PPs of mice fed CE-2 diet, the AIN-93G diet, or AIN-93G diets supplemented with representative CE-2 ingredients for 4 weeks starting from 3 weeks of age. Representative plots of GL-7 and Bcl-6 staining within the CD19$^+$ gate are shown in (B), and quantification of frequency and total number is shown in (C) ($n$ = 6 mice/group, mean ± s.d.).

(D-I) Flow cytometry analysis of lymphocytes in the distal PPs of germ-free (GF) mice fed either the AIN-93G or the AIN-93G + Soy diet for 4 weeks starting from 3 weeks of age. (D and E) CXCR5$^+$Bcl-6$^+$ Tfh cells. Representative plots of CXCR5 and Bcl-6 staining within the CD4$^+$TCRβ$^+$Foxp3$^-$CD25$^-$ gate are shown in (D), and quantification of frequency and total number is shown in (E) ($n$ = 6 mice/group, mean ± s.d.). (F and G) Bcl-6$^+$GL-7$^+$ GCB cells. Representative plots of GL-7 and Bcl-6 staining within the CD19$^+$ gate are shown in (F), and quantification of frequency and total number is shown in (G) ($n$ = 6 mice/group, mean ± s.d.). (H and I) IgA$^+$ B cells. Representative plots of IgA staining within the CD19$^+$ gate are shown in (H), and quantification of frequency and total number is shown in (I) ($n$ =6 mice/group, mean ± s.d.).

Data are representative of at least two independent experiments. Statistical analysis was performed by (C) one-way ANOVA followed by Dunnett's multiple comparisons test, or (E, G, and I) Welch's t-test. *$p < 0.05$; **$p < 0.01$; ***$p < 0.001$.

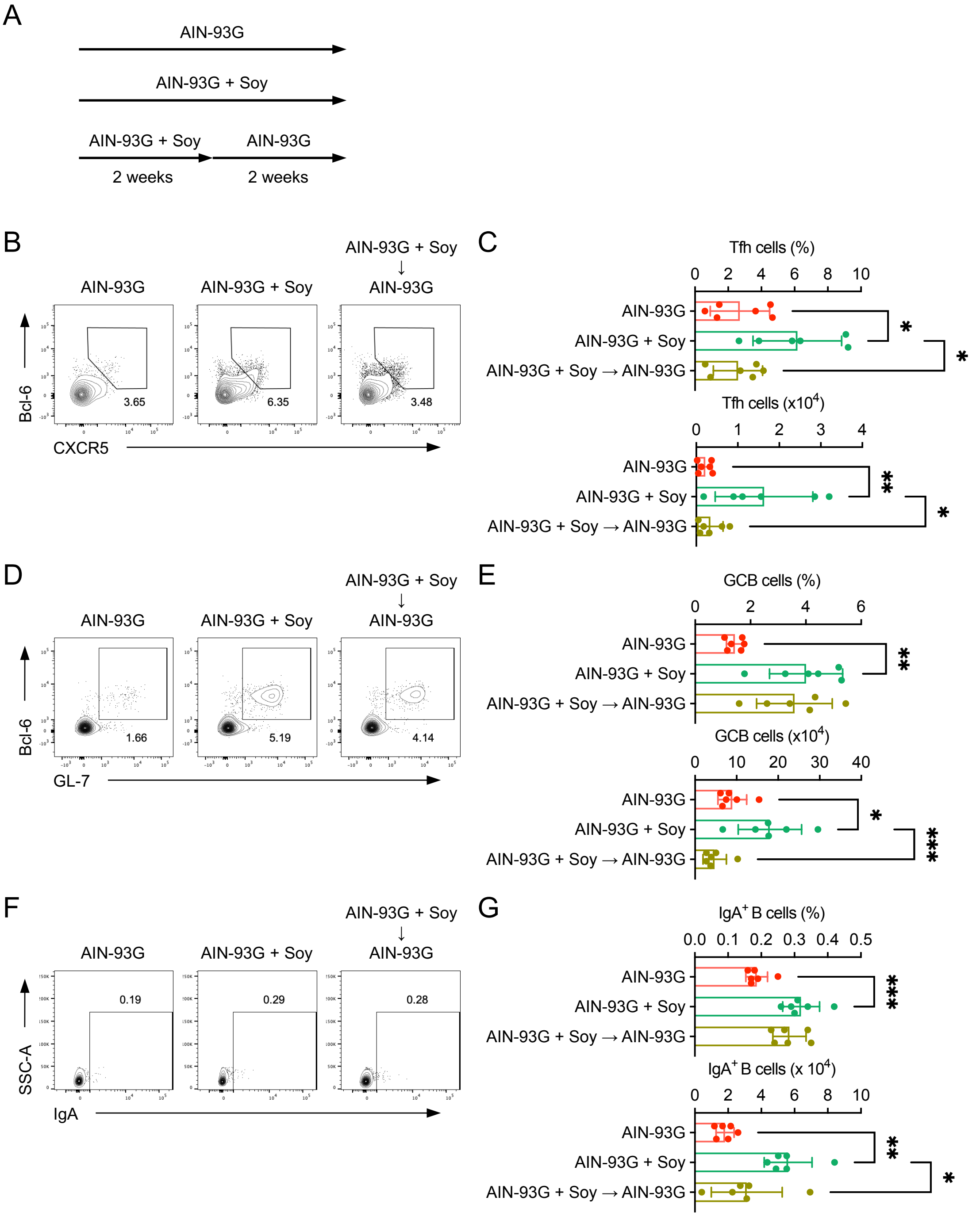


**Figure S3. Continuous intake of soy is essential for maintaining Tfh, GCB, and IgA$^+$ B cells in PPs**

(A) Experimental design for soy intake cessation. Mice were fed the AIN-93G + Soy diet for two weeks and then switched to the AIN-93G diet for the following two weeks. Control groups were fed either AIN-93G or AIN-93G + Soy diets continuously for 4 weeks starting from 3 weeks of age.

(B-G) Flow cytometry analysis of lymphocytes in the distal PPs from mice in the indicated groups. (B and C) CXCR5$^+$Bcl-6$^+$ Tfh cells. Representative plots of CXCR5 and Bcl-6 staining within the CD4$^+$TCRβ$^+$Foxp3$^-$CD25$^-$ gate are shown in (B), and quantification of frequency and total number is shown in (C) (*n* = 6 mice/group, mean ± s.d.). (D and E) Bcl-6$^+$GL-7$^+$ GCB cells. Representative plots of GL-7 and Bcl-6 staining within the CD19$^+$ gate are shown in (D), and quantification of frequency and total number is shown in (E) (*n* = 6 mice/group, mean ± s.d.). (F and G) Representative plots of IgA staining within the CD19$^+$ gate are shown in (F), and quantification of frequency and total number is shown in (G) (*n* =6 mice/group, mean ± s.d.).

Data are representative of at least two independent experiments. Statistical analysis was performed by (C, E and G) one-way ANOVA followed by Tukey's multiple comparisons test. *$p < 0.05$; **$p < 0.01$; ***$p < 0.001$.

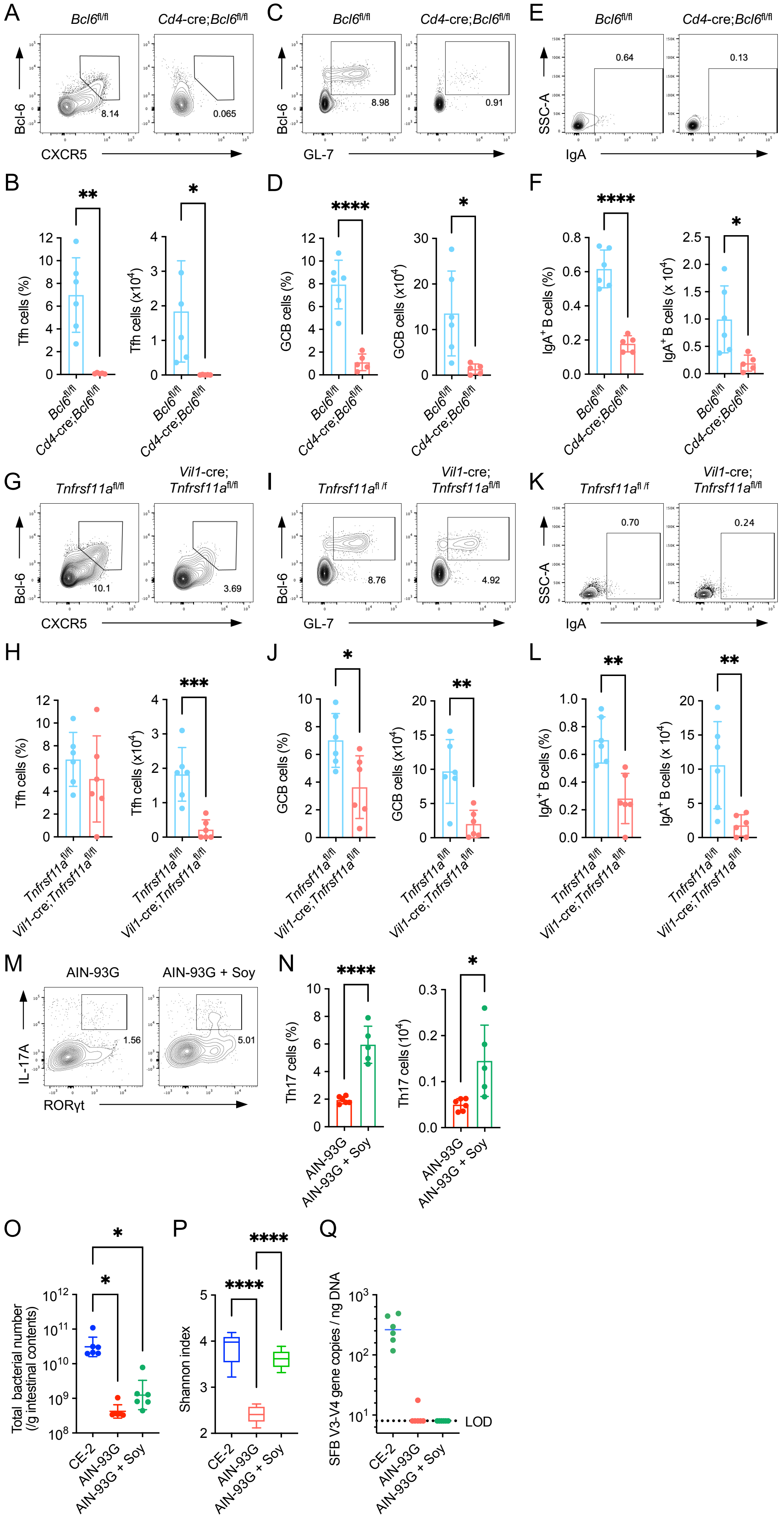
A
Bcl6fl/fl
Cd4-cre;Bcl6fl/fl
Bcl-6
CXCR5
8.14
0.065
B
**
*
Tfh cells (%)
Tfh cells (x10^4)
Bcl6fl/fl
Cd4-cre;Bcl6fl/fl
C
Bcl6fl/fl
Cd4-cre;Bcl6fl/fl
Bcl-6
GL-7
8.98
0.91
D
****
*
GCB cells (%)
GCB cells (x10^4)
E
Bcl6fl/fl
Cd4-cre;Bcl6fl/fl
SSC-A
IgA
0.64
0.13
F
****
*
IgA+ B cells (%)
IgA+ B cells (x 10^4)
G
Tnfrsf11afl/fl
Vil1-cre;
Tnfrsf11afl/fl
Bcl-6
CXCR5
10.1
3.69
H
***
Tfh cells (%)
Tfh cells (x10^4)
Tnfrsf11afl/fl
Vil1-cre;Tnfrsf11afl/fl
I
Tnfrsf11afl /f
Vil1-cre;
Tnfrsf11afl/fl
Bcl-6
GL-7
8.76
4.92
J
*
**
GCB cells (%)
GCB cells (x10^4)
K
Tnfrsf11afl /f
Vil1-cre;
Tnfrsf11afl/fl
SSC-A
IgA
0.70
0.24
L
**
**
IgA+ B cells (%)
IgA+ B cells (x 10^4)
M
AIN-93G
AIN-93G + Soy
IL-17A
RORγt
1.56
5.01
N
****
*
Th17 cells (%)
Th17 cells (10^4)
AIN-93G
AIN-93G + Soy
O
*
*
Total bacterial number
(/g intestinal contents)
CE-2
AIN-93G
AIN-93G + Soy
P
****
****
Shannon index
CE-2
AIN-93G
AIN-93G + Soy
Q
SFB V3-V4 gene copies / ng DNA
LOD
CE-2
AIN-93G
AIN-93G + Soy

**Figure S4. Soy supplementation fails to induce GCB and IgA$^+$ B cells in Tfh cell-deficient mice and M cell-deficient mice**

(A-F) Flow cytometry analysis of Tfh cell-deficient (*Cd4*-Cre;*Bcl6*$^{fl/fl}$) and control (*Bcl6*$^{fl/fl}$) mice fed the AIN-93G + Soy diet for 4 weeks starting from 3 weeks of age. (A and B) CXCR5$^+$Bcl-6$^+$ Tfh cells. Representative plots of CXCR5 and Bcl-6 staining within the CD4$^+$TCRβ$^+$Foxp3$^-$CD25$^-$ gate are shown in (A), and quantification of frequency and total number is shown in (B) ($n$ = 5-6 mice/group, mean ± s.d.). (C and D) Bcl-6$^+$GL-7$^+$ GCB cells. Representative plots of GL-7 and Bcl-6 staining within the CD19$^+$ gate are shown in (C), and quantification of frequency and total number is shown in (D) ($n$ = 5-6 mice/group, mean ± s.d.). (E and F) IgA$^+$ B cells. Representative plots of IgA staining within the CD19$^+$ gate are shown in (E), and quantification of frequency and total number is shown in (F) ($n$ = 5-6 mice/group, mean ± s.d.).

(G-L) Flow cytometry analysis of M cell-deficient (*Vil1*-Cre;*Tnfrsf11a*$^{fl/fl}$) and control (*Tnfrsf11a*$^{fl/fl}$) mice fed the AIN-93G + Soy diet for 4 weeks starting from 3 weeks of age. (G and H) CXCR5$^+$Bcl-6$^+$ Tfh cells. Representative plots of CXCR5 and Bcl-6 staining within the CD4$^+$TCRβ$^+$Foxp3$^-$CD25$^-$ gate are shown in (G), and quantification of frequency and total number is shown in (H) ($n$ = 6 mice/group, mean ± s.d.). (I and J) Bcl-6$^+$GL-7$^+$ GCB cells. Representative plots of GL-7 and Bcl-6 staining within the CD19$^+$ gate are shown in (I), and quantification of frequency and total number is shown in (J) ($n$ = 6 mice/group, mean ± s.d.). (K and L) IgA$^+$ B cells. Representative plots of IgA staining within the CD19$^+$ gate are shown in (K), and quantification of frequency and total number is shown in (L) ($n$ = 6 mice/group, mean ± s.d.).

(M and N) IL-17A$^+$RORγt$^+$ Th17 cells in the distal small intestinal lamina propria of mice fed either the AIN-93G, or AIN-93G + Soy diet for 4 weeks starting from 3 weeks of age. Representative flow cytometry plots of IL-17A and RORγt staining within the CD4$^+$ T cell gate are shown in (M), and quantification of frequency and number of Th17 cells is shown in (N) ($n$ = 6 mice/group, mean ± s.d.).

(O-Q) Analysis of the distal small intestinal microbiota in mice fed the CE-2 or AIN-93G diet for 4 weeks starting from 3 weeks of age. (O) Total bacterial number assessed by qPCR for the 16S rRNA gene ($n$ = 6 mice/group). (P) Species richness (Shannon entropy) (n = 6 mice/group). (Q) SFB copy numbers (n = 6 mice/group) assessed by qPCR (n = 6 mice/group).

Data are representative of at least two independent experiments. Statistical analysis was performed by (B, D, F, H, J, L, and N) Welch's t-test or (O, P, and Q) one-way ANOVA followed by Dunnett's multiple comparisons test. *$p$ < 0.05; **$p$ < 0.01; ***$p$ < 0.001; ****$p$ < 0.0001.

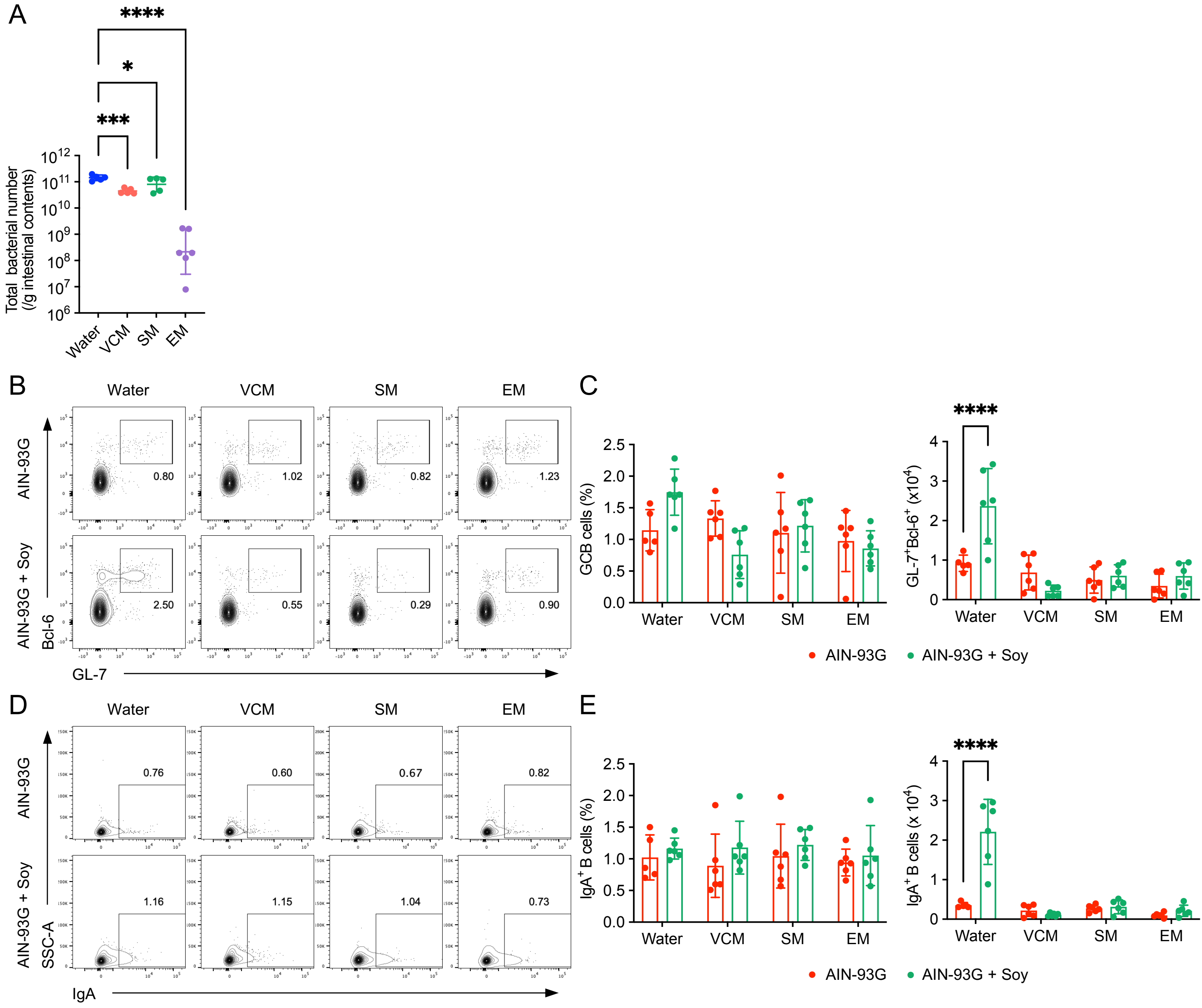


**Figure S5. Antibiotic treatment diminishes soy diet-mediated induction of GCB and IgA⁺ B cells**

(A) Total bacterial number in the distal small intestinal content of mice orally administered with different antibiotics or water (control). Mice fed the AIN-93G + Soy diet and were given drinking water containing vancomycin (VCM; 0.5 g/L), streptomycin (SM; 1 g/L), or erythromycin (EM; 1 g/L) for 4 weeks starting from 3 weeks of age. (*n* = 6 mice/group).

(B-E) Flow cytometry analysis of lymphocytes in the distal PPs of mice orally administered with different antibiotics or water (control). Mice fed either the AIN-93G or the AIN-93G + Soy diet were given drinking water containing vancomycin (VCM; 0.5 g/L), streptomycin (SM; 1 g/L), or erythromycin (EM; 1 g/L) for 4 weeks starting from 3 weeks of age. (*n* = 6 mice/group). (B and C) GL-7⁺Bcl-6⁺ GCB cells. Representative flow cytometry plots of GL-7 and Bcl-6 staining within the CD19⁺ gate are shown in (B), and quantification of frequency and total number is shown in (C) (*n* = 6 mice/group, mean ± s.d.). (D and E) IgA⁺ B cells. Representative flow cytometry plots of IgA staining within the CD19⁺ gate are shown in (D), and quantification of frequency and total number is shown in (E) (*n* = 6 mice/group, mean ± s.d.).

Data are representative of at least two independent experiments. Statistical analysis was performed by (A) one-way ANOVA followed by Dunnett's multiple comparisons test, and by (C and E) two-way ANOVA followed by Šídák's multiple comparisons test. *$p < 0.05$; ***$p < 0.001$; ****$p < 0.0001$.

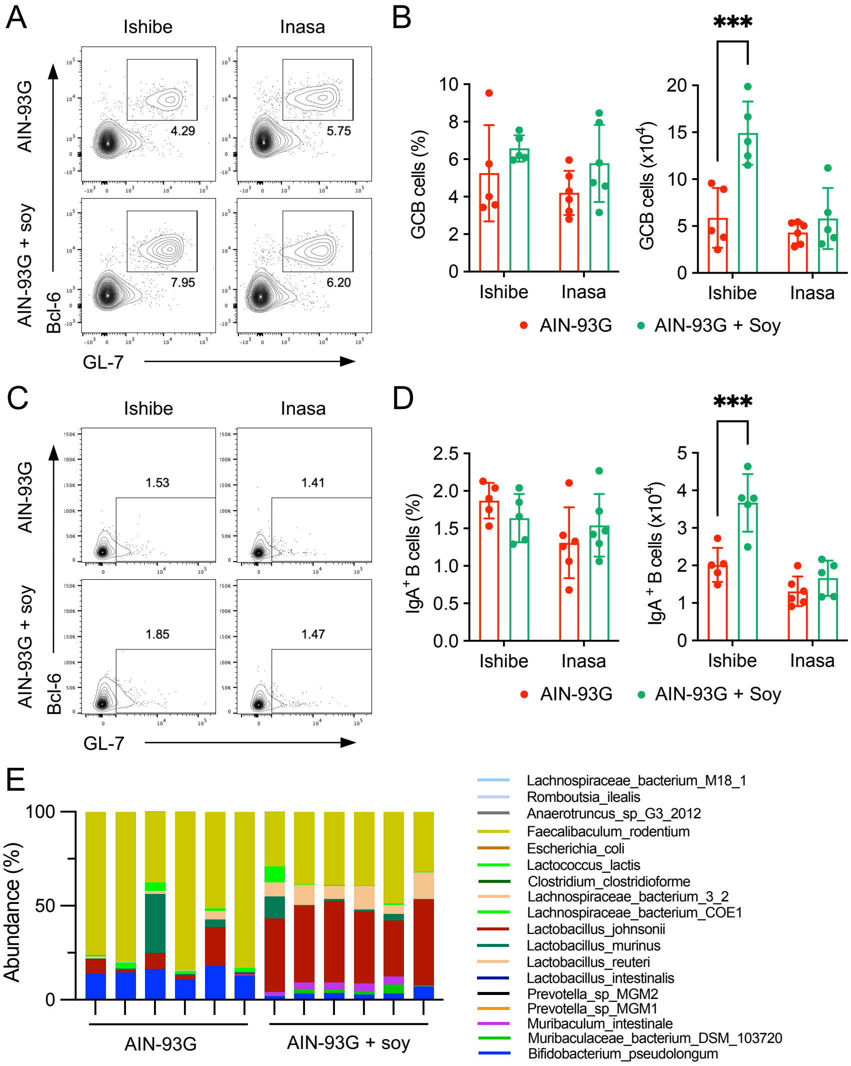


**Figure S6. Soy supplementation fails to induce GCB and IgA⁺ B cells in the distal PPs of mice from the Inasa facility**

(A-D) Flow cytometry analysis of lymphocytes in the distal PPs of mice from two different breeding facilities, Ishibe (CLEA Japan) and Inasa (Japan SLC). Mice were fed either the AIN-93G or the AIN-93G + Soy diet for 4 weeks starting from 3 weeks of age. (A and B) GL-7$^+$Bcl-6$^+$ GCB cells. Representative flow cytometry plots of GL-7 and Bcl-6 staining within the CD19$^+$ gate are shown in (A), and quantification of frequency and total number is shown in (B) ($n$ = 6 mice/group, mean ± s.d.). (C and D) IgA$^+$ B cells. Representative flow cytometry plots of IgA staining within the CD19$^+$ gate are shown in (C), and quantification of frequency and total number is shown in (D) ($n$ = 6 mice/group, mean ± s.d.).

(E) Shotgun metagenome sequencing analysis of the distal small intestinal microbiota from Ishibe facility mice fed either the AIN-93G or the AIN-93G + Soy diet. The composition is shown at the species level.

Data in (A-D) are representative of at least two independent experiments. Statistical analysis was performed by (B and D) two-way ANOVA followed by Šídák's multiple comparisons test. ***$p < 0.001$.

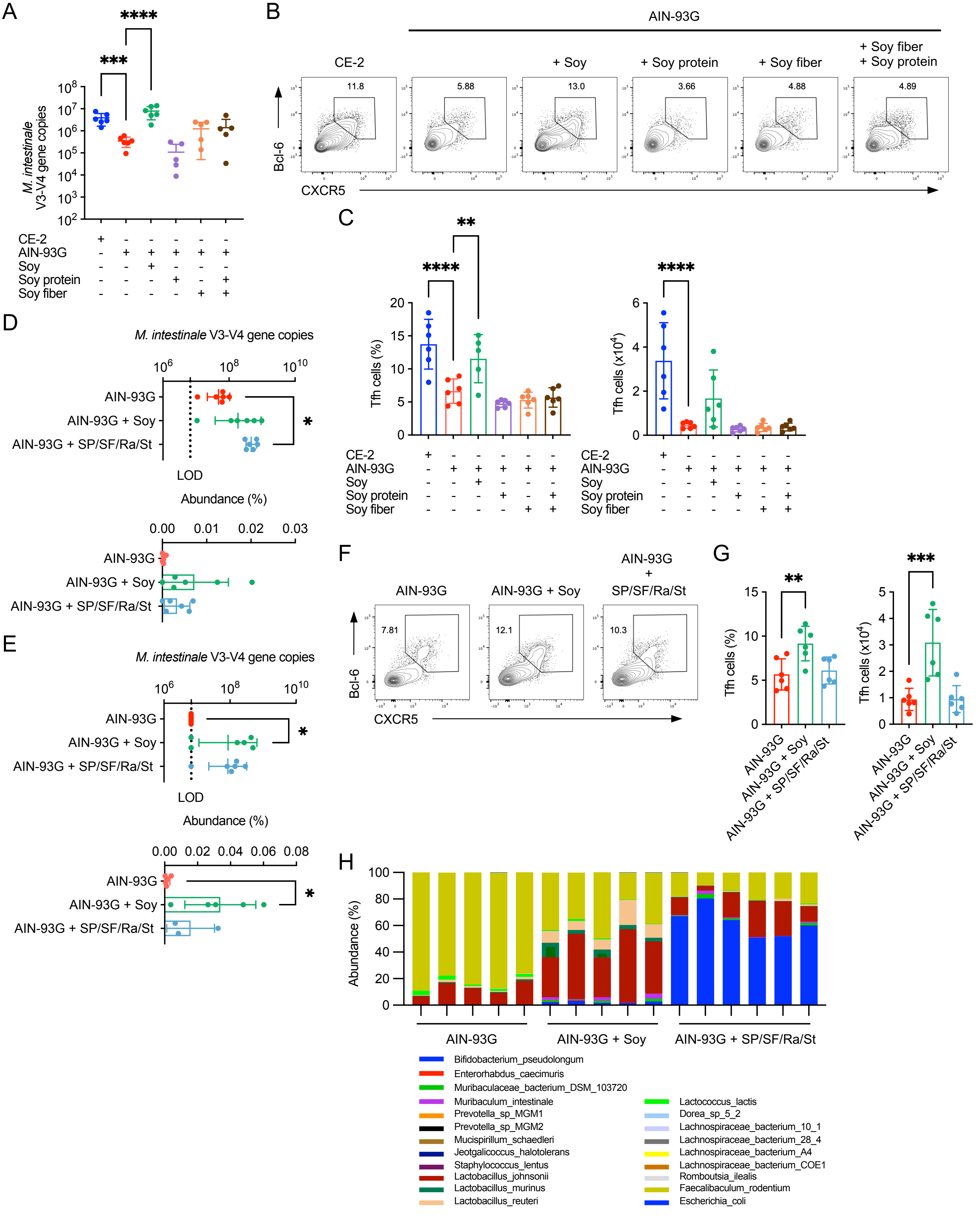


**Figure S7. Soy oligosaccharides are required for the soy-mediated growth of *M. intestinale***

(A-C) Effects of soy protein and fiber supplementation. Mice were fed the indicated diets for 4 weeks starting from 3 weeks of age. (A) The number of *M. intestinale* in the distal small intestinal content, as assessed by qPCR ($n$ = 6 mice/group). (B and C) CXCR5$^+$Bcl-6$^+$ Tfh cells in the distal PPs. Representative flow cytometry plots of CXCR5 and Bcl-6 staining within the CD4$^+$TCRβ$^+$Foxp3$^-$CD25$^-$ gate are shown in (B), and quantification of frequency and total number is shown in (C) ($n$ = 6 mice/group, mean ± s.d.).

(D-H) Effects of soy oligosaccharide supplementation. Mice were fed the indicated diets for 4 weeks starting from 3 weeks of age. The AIN-93G + SP/SF/Ra/St diet was supplemented with soy protein, soy fiber, 2% raffinose, and 1% stachyose. (D and E) The number (by qPCR) and relative abundance (by 16S rRNA gene sequencing) of *M. intestinale* in the distal small intestinal content (D) and mucus layer (E) ($n$ = 6 mice/group). (F and G) CXCR5$^+$Bcl-6$^+$ Tfh cells in the distal PPs. Representative flow cytometry plots of CXCR5 and Bcl-6 staining within the CD4$^+$TCRβ$^+$Foxp3$^-$CD25$^-$ gate are shown in (F), and quantification of frequency and total number is shown in (G) ($n$ = 6 mice/group, mean ± s.d.). (H) Shotgun metagenome sequencing analysis of the distal small intestinal microbiota, shown at the species level (n = 6 mice/group).

Data are representative of at least two independent experiments. Statistical analysis was performed by (A, C, E, and G) one-way ANOVA followed by Tukey's multiple comparisons test. *$p < 0.05$; **$p < 0.01$; ***$p < 0.001$; ****$p < 0.0001$.

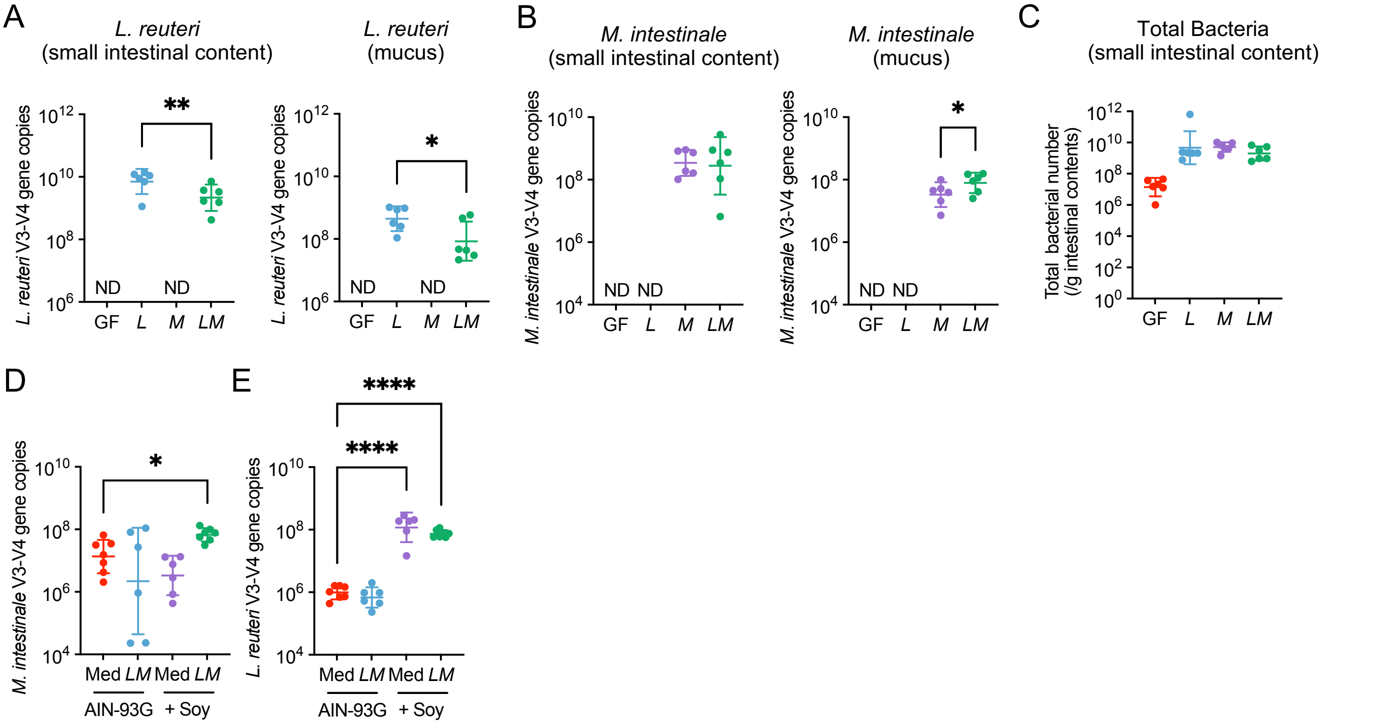


**Figure S8. Successful colonization of *L. reuteri* and *M. intestinale* in gnotobiotic and Inasa mice**

(A-C) Bacterial colonization in gnotobiotic mice. Germ-free (GF) mice were mono-colonized with *L. reuteri* (*L*) or *M. intestinale* (*M*), or co-colonized with both (*L. reuteri* and *M. intestinale*; *LM*). All mice were fed the AIN-93G + Soy diet. (A and B) The number of *L. reuteri* (A) and *M. intestinale* (B) in the distal small intestinal content and mucus layer, as quantified by qPCR using species-specific primers ($n$ = 6 mice/group, mean ± s.d.). ND, not detected. (C) Total bacterial number in the distal small intestinal content, as quantified by qPCR using universal 16S rRNA primers ($n$ = 6 mice/group, mean ± s.d.).

(D and E) Bacterial colonization in conventionalized mice from the Inasa facility. Mice were inoculated with a control culture medium (Med) or a mixture of *L. reuteri* and *M. intestinale* (*LM*), and fed either the AIN-93G diet or the AIN-93G + Soy diet. The number of *L. reuteri* (D) and *M. intestinale* (E) in the distal small intestinal content, as quantified by qPCR (n = 6 mice/group, mean ± s.d.).

Data are representative of at least two independent experiments. Statistical analysis was performed by (A-C) one-way ANOVA followed by Tukey's multiple comparisons test, or by (D and E) two-way ANOVA followed by Šídák's multiple comparisons test. *$p$ < 0.05; **$p$ < 0.01; ****$p$ < 0.0001.

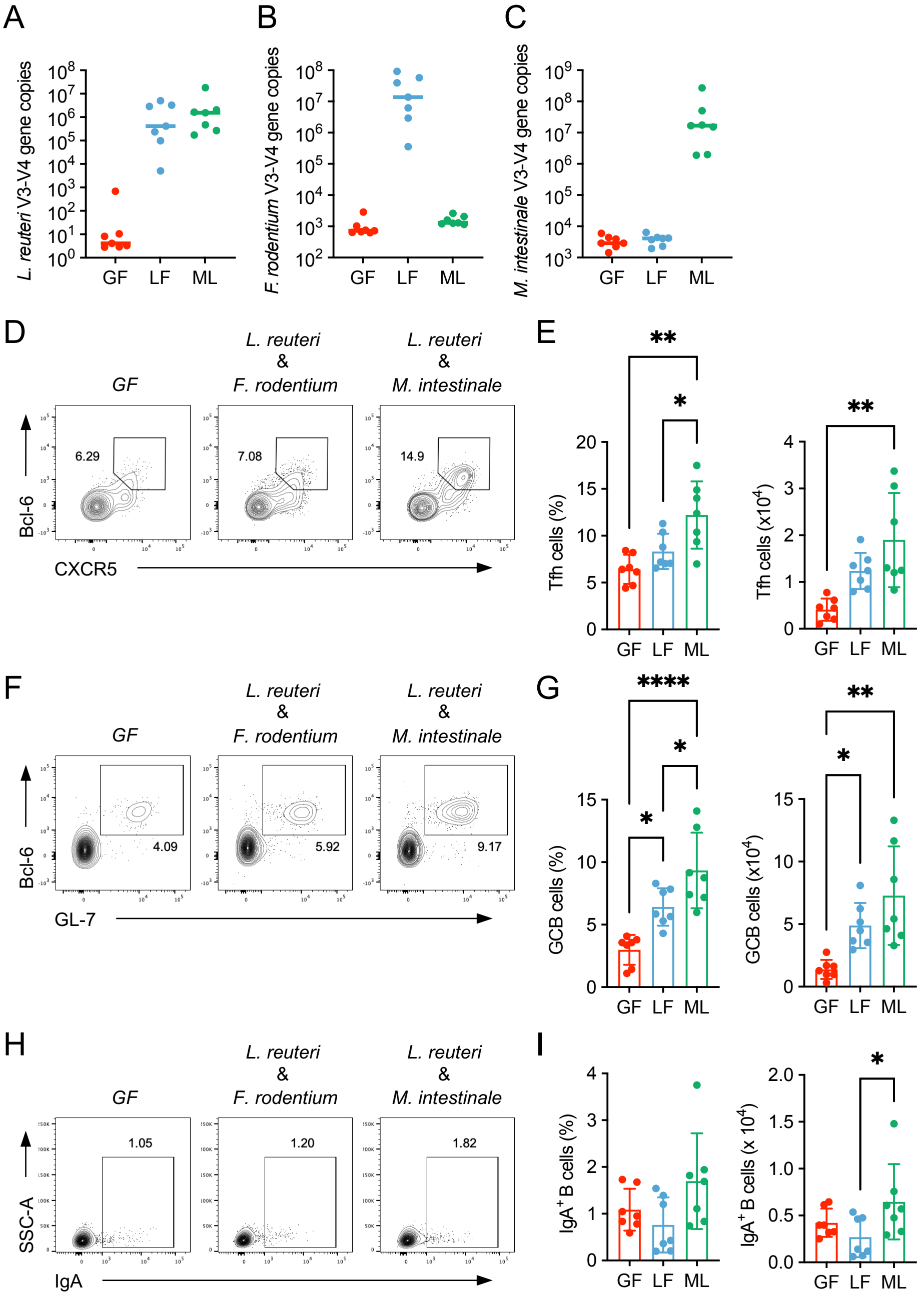


**Figure S9. Co-colonization with *L. reuteri* and *F. rodentium* fails to efficiently induce PP Tfh, GCB, and IgA⁺ B cells**

(A-C) Bacterial colonization in the distal small intestinal content of gnotobiotic mice. Groups include germ-free (GF), mice co-colonized with *L. reuteri* and *F. rodentium* (LF), and mice co-colonized with *L. reuteri* and M. intestinale (LM). The number of *L. reuteri* (A), *F. rodentium* (B), and *M. intestinale* (C) was quantified by qPCR using species-specific primers ($n$ = 7 mice/group, mean ± s.d.).

(D-I) Flow cytometry analysis of lymphocytes in the distal PPs of the gnotobiotic mouse groups described above. (D and E) $CXCR5^+Bcl\text{-}6^+$ Tfh cells. Representative flow cytometry plots of CXCR5 and Bcl-6 staining within the $CD4^+TCR\beta^+Foxp3^-CD25^-$ gate are shown in (D), and quantification of frequency and total number is shown in (E) ($n$ = 7 mice/group, mean ± s.d.). (F and G) $GL\text{-}7^+Bcl\text{-}6^+$ GCB cells. Representative flow cytometry plots of GL-7 and Bcl-6 staining within the $CD19^+$ gate are shown in (F), and quantification of frequency and total number is shown in (G) ($n$ = 6-7 mice/group, mean ± s.d.). (H and I) $IgA^+$ B cells. Representative flow cytometry plots of IgA staining within the $CD19^+$ gate are shown in (H), and quantification of frequency and total number is shown in (I) ($n$ = 7 mice/group, mean ± s.d.).

Data are representative of at least two independent experiments. Statistical analysis was performed by (E, G, and I) one-way ANOVA followed by Tukey's multiple comparisons test. *$p < 0.05$; **$p < 0.01$; ****$p < 0.0001$.

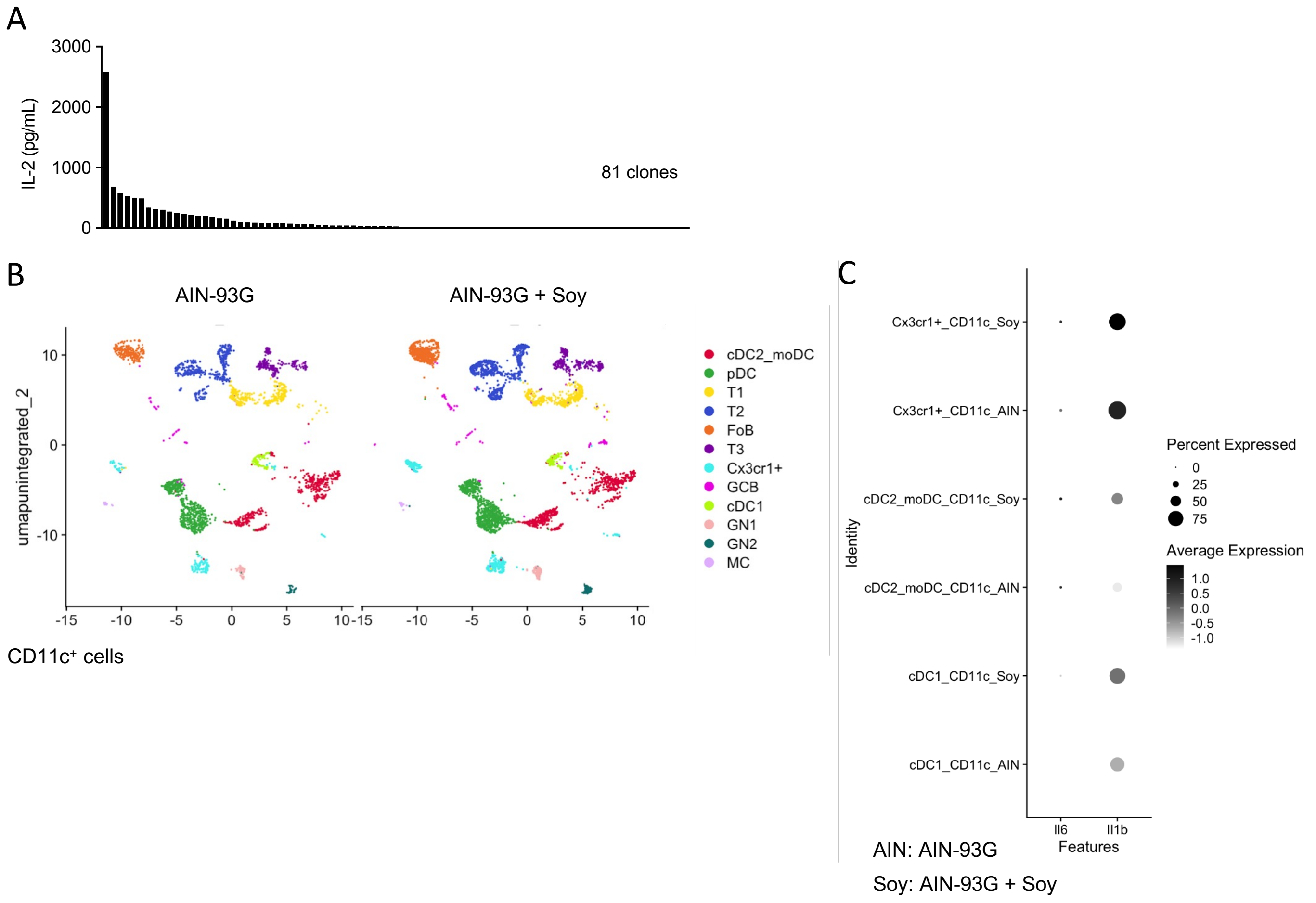


**Figure S10. *L. reuteri* provides antigens for PP Tfh cells, and soy supplementation induces IL-1β from myeloid cell subsets in distal PPs**

(A) IL-2 concentration in the supernatant of 81 unique PP Tfh hybridoma clones. Each clone, possessing a unique TCRα and β chain, was co-cultured for 24 hours with BMDCs in the presence of *L. reuteri* lysates. Data are from two technical replicates.

(B) Uniform Manifold Approximation and Projection (UMAP) plot of $CD11c^+$ cells isolated from the distal PPs of mice fed either the AIN-93G diet (labeled as AIN) or the AIN-93G + Soy (labeled as Soy) diet.

(C) Dot plot showing the expression of *Il1b* and *Il6* genes across myeloid cell clusters identified in (B), including conventional dendritic cell 1 (cDC1), cDC2, monocyte-derived DCs (moDC), and $CX3CR1^+$ monocytes.

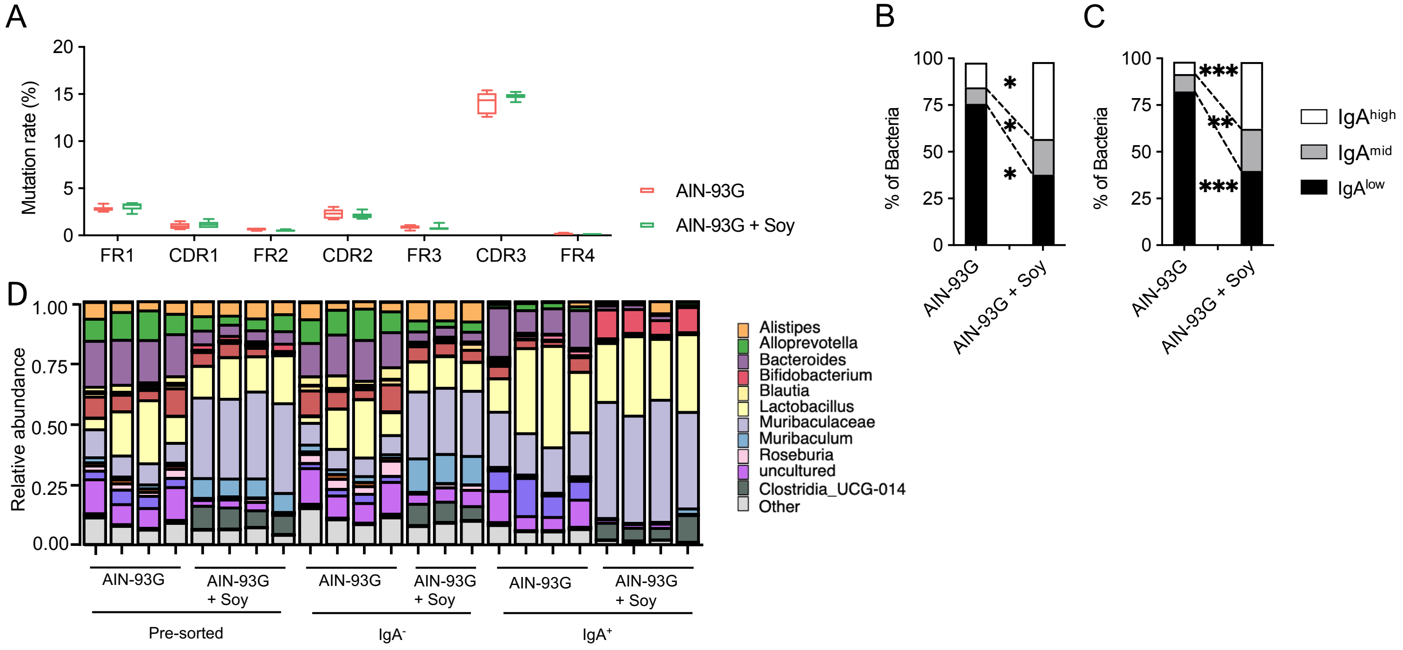


**Figure S11. Composition of IgA-coated bacteria and IgH mutation analysis**

(A) Mutation rates of the immunoglobulin heavy chain (IgH) variable region from IgA-producing plasma cells. Tissue fragments were isolated from the distal small intestine of mice fed either the AIN-93G diet or the AIN-93G + Soy diet. Mutation rates at each complementarity-determining region (CDR1-3) and framework region (FWR1-3) were calculated from IgH sequencing results (*n* = 6 mice/group, mean ± s.d.).

(B and C) Flow cytometry analysis of IgA-coated bacteria in proximal (B) and distal (C) small intestinal content from mice fed either the AIN-93G or the AIN-93G + Soy diet. (n = 6 mice/group).

(D) Genus-level composition of IgA-coated and IgA-unbound bacteria from fecal samples of mice fed either the AIN-93G diet or the AIN-93G + Soy diet. Bacterial fractions were sorted by FACS based on IgA binding and analyzed by 16S rRNA gene sequencing. The composition of the total, unsorted fecal microbiota (Pre-sorted) is also shown for comparison (*n* = 3-4 mice/group).

Data are representative of at least two independent experiments. Statistical analysis was performed by (A) two-way ANOVA followed by Šídák's multiple comparisons test, and by (B) Welch's t-test or Student's t-test. *$p < 0.05$.

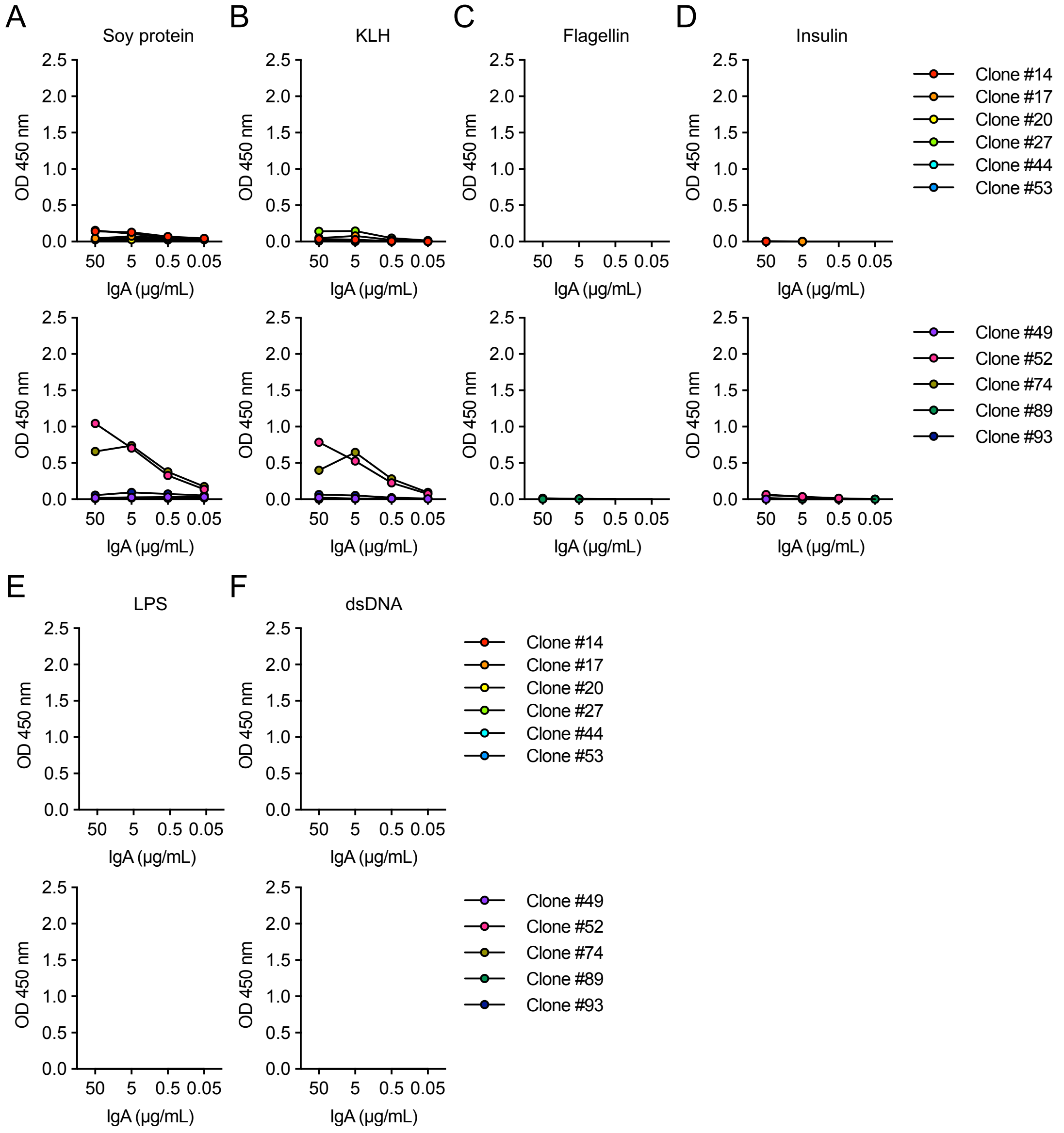


**Figure S12. Reactivity profile of monoclonal IgA hybridoma clones**

Binding of IgA-producing hybridoma culture supernatants to various antigens was assessed by ELISA. The tested antigens include (A) Soy protein, (B) Keyhole limpet hemocyanin (KLH), (C) Flagellin, (D) Insulin, (E) Lipopolysaccharide (LPS), and (F) double-stranded DNA (dsDNA). See also Table S4 for details of each IgA clone.

**Table S1 Composition of the CE-2 and AIN-93G diets**

| | CE-2 | AIN-93G |
|---|---|---|
| Protein source | whitefish meal, soybean meal, yeast | casein |
| Fat source | cereal germ, soybean oil | soybean oil |
| Fiber source | wheat bran, defatted rice bran, alfalfa meal | cellulose |
| Carbohydreate source | wheat flour, corn, milo | cornstarch, α-cornstarch |
| Vitamins | Vita.A, D3, E, B1, B2, B6, B12, C, niacin, pantothenic acid, biotin, folic acid,choline chloride, inositol | Vitamin Mix (V10037) |
| Minerals | calicum carbonate, salt, ferrous sulfate, manganese sulfate, cobalt sulfate, calcium iodate | Mineral Mix (S10022G) |

**Table S2 Composition of the AIN-93G and AIN-93G + soy diet**

| % | AIN-93G | AIN-93G + 33% Soy |
|---|---|---|
| Casein | 20 | 4.8 |
| L-cystine | 0.3 | 0.3 |
| Cornstarch | 34.7486 | 27.6 |
| α-cornstarch | 13.2 | 13.2 |
| Sucrose | 10 | 10 |
| Soy oil | 7 | 0 |
| Cellose powder | 5 | 0 |
| AIN-93G mineral mix | 3.5 | 3.5 |
| AIN-93G vitamin mix | 6 | 6 |
| Choline bitartrate | 0.25 | 0.25 |
| Tertiary butyl hydroquinone | 0.0014 | 0.0014 |
| Soy powder | 0 | 34.3 |
| total | 100 | 99.9514 |

**Table S3 Composition of AIN-93G and modified AIN-93G**

| % | AIN-93G | AIN-93G (Soy protein) | AIN-93G (Soy fiber) | AIN-93G (Soy protein & fiber) | AIN-93G (Soy protein & fiber + raffinose + stachyose) |
|---|---|---|---|---|---|
| casein | 20 | 0 | 20 | 0 | 0 |
| L-cystine | 0.3 | 0.3 | 0.3 | 0.3 | 0.3 |
| cornstarch | 34.7486 | 34.7486 | 27.6 | 27.6 | 27.6 |
| α-cornstarch | 13.2 | 13.2 | 13.2 | 13.2 | 13.2 |
| sucrose | 10 | 10 | 10 | 10 | 10 |
| soy oil | 7 | 7 | 7 | 7 | 7 |
| cellose powder | 5 | 5 | 0 | 0 | 0 |
| AIN-93G mineral mix | 3.5 | 3.5 | 3.5 | 3.5 | 3.5 |
| AIN-93G vitamin mix | 6 | 6 | 6 | 6 | 6 |
| Choline bitartrate | 0.25 | 0.25 | 0.25 | 0.25 | 0.25 |
| tertiary butyl hydroquinone | 0.0014 | 0.0014 | 0.0014 | 0.0014 | 0.0014 |
| soy protein | | 20 | 0 | 20 | 20 |
| soy fiber | | 0 | 12 | 12 | 12 |
| raffinose | | 0 | 0 | 0 | 1 |
| stachyose | | 0 | 0 | 0 | 2 |
| Total | 100 | 99.75 | 99.6014 | 99.6014 | 102.6014 |

**Table S4 Molecular characteristics of IgA hybridoma heavy chains**

| Clone ID | V Gene Allele | D Gene Allele(s) | J Gene Allele | V Gene Mutations (n) | V Gene Mutation Rate (%) |
|---|---|---|---|---|---|
| 14 | IGHV1-80*00 | IGHD2-700 / IGHD5-200 | IGHJ2*00 | 21 | 7.84 |
| 17 | IGHV1-80*00 | IGHD1-1*00 | IGHJ2*00 | 5 | 1.85 |
| 20 | IGHV3-6*00 | IGHD2-300 / IGHD2-700 | IGHJ1*00 | 10 | 3.70 |
| 27 | IGHV9-3*00 | IGHD1-1*00 | IGHJ2*00 | 5 | 1.85 |
| 44 | IGHV6-3*00 | IGHD4-1*00 | IGHJ1*00 | 14 | 5.19 |
| 49 | IGHV3-5*00 | IGHD4-1*00 | IGHJ2*00 | 1 | 0.37 |
| 52 | IGHV2-2*00 | IGHD4-1*00 | IGHJ3*00 | 6 | 2.22 |
| 53 | Not determined | Not determined | Not determined | Not determined | Not determined |
| 74 | IGHV1-80*00 | IGHD2-8*00 | IGHJ4*00 | 8 | 2.96 |
| 89 | IGHV7-1*00 | Not identified | IGHJ2*00 | 1 | 0.37 |
| 93 | IGHV9-3*00 | IGHD2-300 / IGHD2-800 | IGHJ2*00 | 10 | 3.70 |

**Table S5 Primers and probes used for qPCR**

| Target | Type | Sequence (5' → 3') |
|---|---|---|
| SFB | Forward Primer | TGAGCGGAGATATATGGAGC |
| | Probe | /56-FAM/ACTTAGCAG/ZEN/CGAACGGGTGAGTAACA/3IABkFQ/ |
| | Reverse Primer | CATGCAACTATATAGCTATATGCGG |
| *L. reuteri* | Forward Primer | ACCGAGAACACCGCGTTATTT |
| | Probe | /56-FAM/ATCGCTAAC/ZEN/TCAATTAAT/3IABkFQ/ |
| | Reverse Primer | CATAACTTAACCTAAACAATCAAAGATTGTCT |
| *M. intestinale* | Forward Primer | TCAAGTCAGCGGTAAAAATTCG |
| | Probe | /56-FAM/CAACCCCGT/ZEN/CGTGCC/3IABkFQ/ |
| | Reverse Primer | CCCACTCAAGAACATCAGTTTCAA |
| *F. rodentium* | Forward Primer | CCGGGAATACGCTCTGGAAA |
| | Reverse Primer | GCCAACCAACTAATGCACCG |
